# 22kHz Ultrasonic Vocalizations and Body Freezing May Represent Two Systems of Fear in Rodent Fear Conditioning Paradigms

**DOI:** 10.1101/2023.06.02.543520

**Authors:** Benhuiyuan Zheng, Lili Bao, Delin Yu, Bin Yin

## Abstract

The “two-system” framework of fear proposed by LeDoux and Pine (2016) sparked a discussion on the understanding of fear, prompting a reevaluation of rodent fear conditioning studies. We propose that 22kHz ultrasonic vocalizations (USV) may symbolize the subjective negative emotional states in rats. To evaluate this, we designed a series of fear conditioning experiments with varied parameters, comparing the expression of 22kHz USV and body freezing – a traditional fear index. These expressions were further assessed in fear generalization tests. Our findings suggest a distinct separation between body freezing and 22kHz vocalizations in fear conditioning under different conditions. The results indicate that 22kHz USV may more accurately signify the subjective state of fear, whereas body freezing may denote an automatic defensive response in rats. Consequently, we posit that the two-system fear framework may extend to rodent fear conditioning paradigms. Therefore, researchers should place greater emphasis on 22kHz vocalizations when investigating the subjective state of fear.

## Introduction

Fear, as a potent negative emotion, mirrors an individual’s reaction to perceived threats. Its study not only elucidates the dynamic interplay between individuals and their environment, but also enriches our understanding of the mechanisms and potential treatments for fear-related psychiatric disorders such as phobias and post-traumatic stress disorder (PTSD) [1–3]. Consequently, a clear definition of fear serves as a cornerstone in fear research.

Traditionally, fear is perceived as an amalgamated signal that initiates sensory, behavioral, and physiological changes in an individual upon encountering a threat [4]. Thus, an individual’s fear acquisition is typically inferred from observable fear-related behaviors. However, LeDoux and Pine introduced a novel “two-system” framework, redefining fear as a subjective feeling, and the behavioral and physiological changes as defensive responses facilitated by separate yet interrelated neural circuits [5]. This proposal stimulated an active discourse about the concept of fear, prompting researchers to reevaluate how fear can be thoroughly characterized.

Rodent models of fear, specifically the Pavlovian fear conditioning paradigm, are widely employed. In this model, subjects learn to correlate a neutral stimulus (Conditioned Stimulus, CS) with an aversive stimulus (Unconditioned Stimulus, US) following successful pairings. This paradigm has elucidated intricate behavioral and neurophysiological mechanisms underpinning fear for decades [6–8]. Generally, fear responses are gauged through body freezing – a state of total immobility and tension due to muscular tonicity, widely accepted as a standard indicator of fear in rodents [9, 10] . However, LeDoux and Pine’s two-system fear framework posits that freezing merely reflects a defensive response, not fear itself. Critics argue that, given animals’ inability to express feelings, we cannot experimentally confirm their experience of fear, potentially undermining the validity of animal models. We contend that the challenge lies in measuring subjective fear, not in the theory itself. In this regard, ultrasonic vocalizations (USV) may offer insight into rodents’ subjective emotional states [11]. Our focus is on 22kHz USV, typically elicited in adverse contexts such as imminent predator attacks, pain, social defeat, or post-coital refractory periods [11, 12]. Existing literature suggests a homology between rat 22kHz USV and human crying, with considerable overlap in characteristics, functions, and underlying neural circuits [13].

While 22kHz USV are acknowledged as fear indicators in fear conditioning [14–16], distinct patterns between 22kHz USV and freezing have been observed in rats across different strains, sexes, and tone tests [17], as well as in lesion experiments involving varied brain regions [18, 19]. If fear originates from a unified central system framework, then 22kHz USV and freezing should exhibit similar trends as fear indicators across contexts. However, whether the two kinds of fear expression in rodents stem from a unified response system or multiple independent fear representation systems remains elusive.

To provide further insights into this question, we conducted a series of tests with varying CS and US parameters, drawing from the Pavlovian fear conditioning paradigm. Studies have shown that increasing US intensity or duration yields more overt behavioral responses and robust fear memories [20–22], leading us to investigate the effects of different shock intensities on 22kHz vocalization and freezing expression. Additionally, we manipulated the CS duration during fear learning to examine its relationship with 22kHz USV and freezing, as research has indicated higher freezing levels following shorter CS duration [23]. Similarly, we altered the CS duration during the cue test with the same fear learning experiences. Lastly, we incorporated three rounds of fear generalization tests and a cue retest for each group, as fear generalization represents an adaptive decision to minimize risk and reflects an individual’s fear level towards a threat [24]. In total, we used six experimental groups with diverse CS and US parameters, a standard experimental (CS+) group, and a control (CS-) group.

Our aim is to illuminate the complex relationship between 22kHz USV and body freezing in the context of fear conditioning. In order to do so, we utilized the mean value every five seconds to plot the time series in all experimental phases. This approach offers a novel perspective on previous studies that typically use the average value of the entire trial in rodent fear conditioning. The functionality of fear can be understood within the context of the continuous threat situation of the predatory imminence continuum (PIC) [25]. Therefore, by measuring external fear indicators on a smaller time scale, we can gain a clearer understanding of how dynamic fear-related behavior organizes to confront imminent threats, which is essential in elucidating the relationship between freezing and 22kHz calling in fear. By comprehensively examining these two fear indices under varied conditioning parameters, we hope to advance our understanding of the subjective state of fear in rodents and shed new light on fear research methodologies. We also aim to bridge the gap between traditional and novel conceptualizations of fear, paving the way for more nuanced and effective treatments for fear-related psychiatric disorders.

## Materials and Methods

### Animals

We utilized 128 male Sprague-Dawley rats, weighing between 300-350g, procured from the Experimental Animal Center of Hangzhou Medical College, Hangzhou, Zhejiang, China. The rats were randomly assigned to eight groups and housed in groups of three per cage, except for one cage which housed only two rats. They were maintained under a 12-hour light/dark cycle (lights on at 8:00 PM and off at 8:00 AM) in standard cages, with ad libitum access to food and water. All subjects had restraint experiences before they went into the main experiments in this study as previous studies have shown that such experiences may enhance freezing behavior in response to salient CS [26, 27]. All procedures were approved by *the Institutional Experimental Animal Care and Use Committee of Fujian Normal University* (protocol number: IACUC-20210041), strictly abided by the ARRIVE 2.0 guidelines [28] and in accordance with the National Research Council’s *Guide for the Care and Use of Laboratory Animals*.

### Apparatus and Experiment Procedures

The study employed a fear conditioning system (Coulbourn Instruments, Holliston, MA), comprising two identical chambers (17.75cm * 17.75cm * 30.5cm) made of aluminum and Plexiglas walls with a stainless-steel rod flooring connected to a shock generator (Coulbourn Instruments). To prevent interference between the two chambers during USV collection, they were vertically arranged and separated by a foam box, and each chamber was enclosed within a sound-insulating cubicle.

Rats underwent fear conditioning in these modular chambers. Contexts for habitation, learning, and contextual tests were cleaned with 75% alcohol (Context A). For the cue test and generalization tests, the stainless-steel floor was swapped with an acrylic *Lego* plate, wrapped with black tape, and cleaned with 3% acetic acid (Context B). Stimulus presentation and video recording were automated using FreezeFrame5 software (Coulbourn Instruments).

The comprehensive experimental procedure along with the corresponding parameters are outlined in **Table 1** and diagramed in **Figure 1**. On the first day, each rat was given a 10-minute habituation period to freely explore the fear conditioning chamber. On the second day, all groups, excluding the control group, underwent three trials of conditioned stimulus (CS) – unconditioned stimulus (US) pairings following a 10-minute baseline period. The standard group underwent three pairs of standard fear learning, with the CS set as a 20 s, 83-85 dB, 9000 Hz pure tone, the US set as a 1 s, 0.7 mA footshock that ended concurrently with the CS, and the inter-trial interval (ITI) set as 180 s. These parameters were in accordance with previously established protocols in LeDoux’s [29] and our labs [30]. The weak fear conditioning (wFC) and strong fear conditioning (sFC) groups received a weaker (0.5 mA, 0.5 s) and a stronger (0.9 mA, 2 s) footshock respectively, while the CS duration remained consistent. The short test (ST) and long test (LT) groups underwent the same standard fear learning process, whereas the short learning (SL) and long learning (LL) groups were exposed to shorter (5 s) and longer (80 s) CS durations respectively, with a standard US. The control group, meanwhile, only received the CS without the US. On the third day, all groups were put through a 3 min contextual test followed by a 9 min cue test. The baseline duration for the cue test was set at 180 s, including two CS presentation trials with a 180 s ITI. With the exception of the ST and LT groups which had shorter (5 s) and longer (80 s) CS durations respectively, all groups underwent a 20s CS exposure each trial. On the fourth and fifth days, all groups underwent three rounds of generalization tests (the CS was set to a 11840 Hz tone, a 7000 Hz tone, and white noise, respectively), followed by a retest with the original CS. The procedural layout of all generalization tests and retests was identical to that of the cue test. The rats were removed from the chamber at the end of the final trial for each test during the experiment.

**Figure 1.**
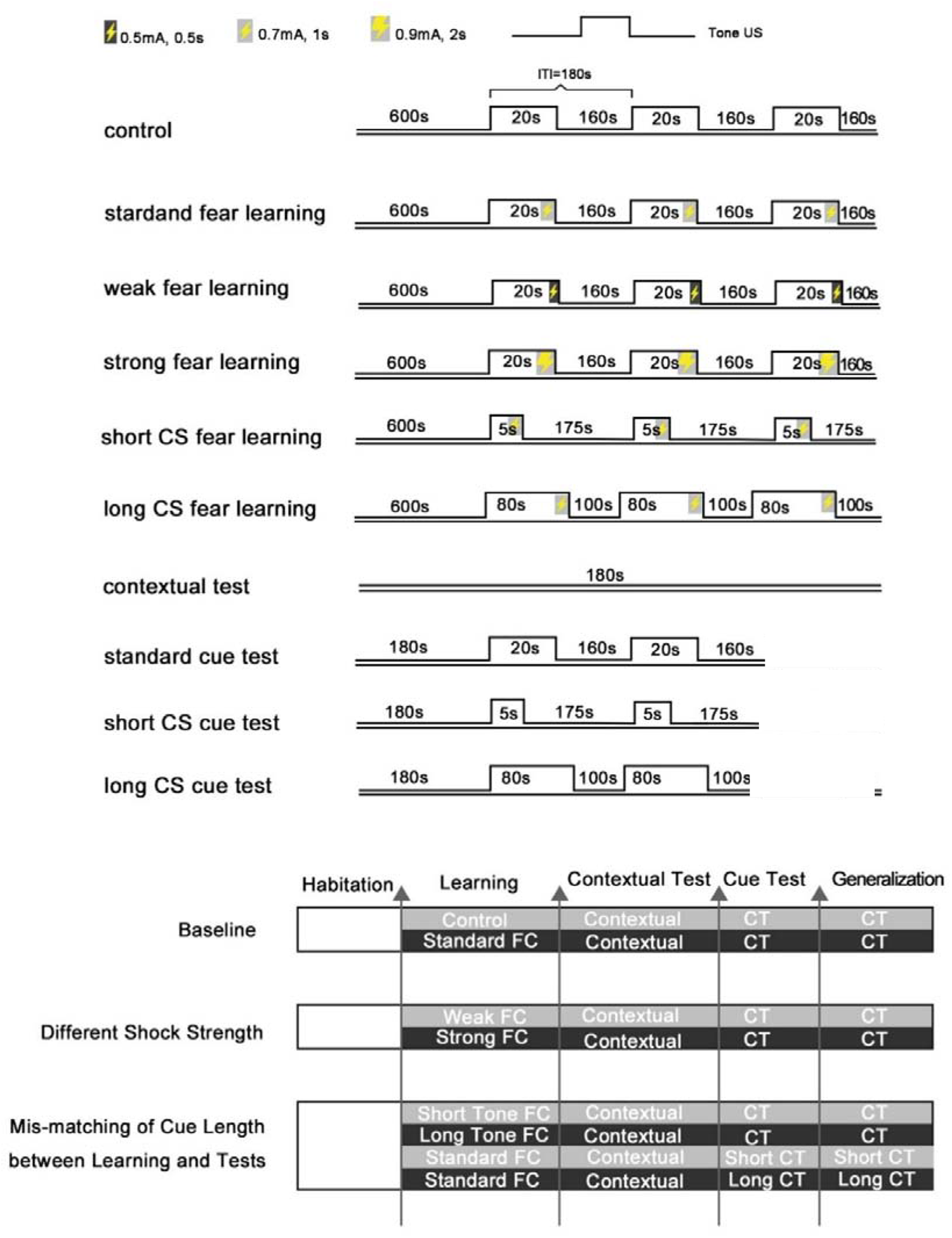
Schematic Overview of the Experimental Design. The study comprised eight groups: (1) Control group, which underwent fear learning without footshock and a standard cue test, (2) Standard group, which underwent standard fear learning and a standard cue test, (3) Weak Fear Conditioning (wFC) group, which underwent weak fear learning and a standard cue test, (4) Strong Fear Conditioning (sFC) group, which underwent strong fear learning and a standard cue test, (5) Short Learning (SL) group, which underwent short CS fear learning and a standard cue test, (6) Long Learning (LL) group, which underwent long CS fear learning and a standard cue test, (7) Short Test (ST) group, which underwent standard fear learning and short CS cue test, and (8) Long Test (LT) group, which underwent standard fear learning and long CS cue test. All groups followed an identical sequence of experimental phases: habituation, fear learning, contextual test, cue test, and fear generalization tests

**Table 1.**
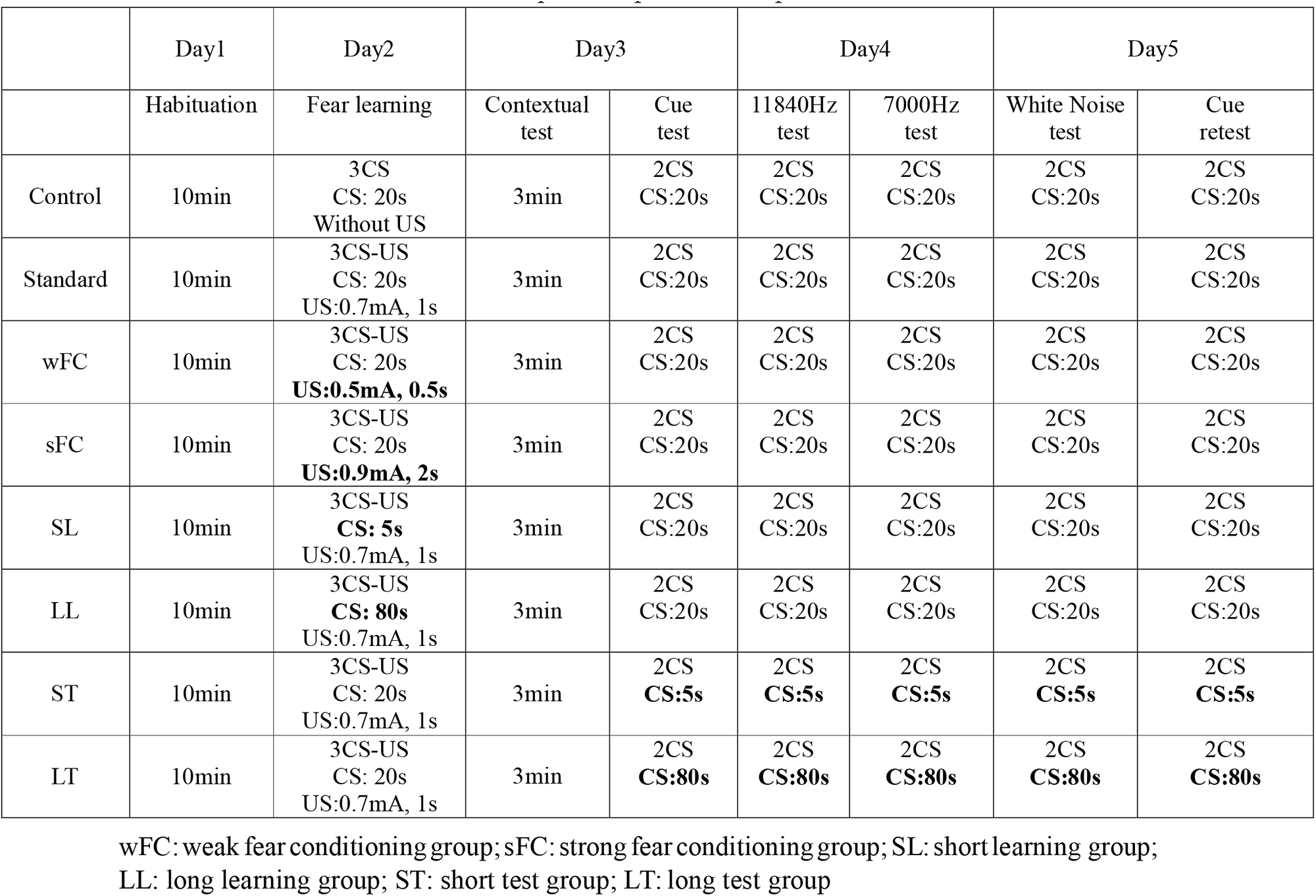
Experiment protocols and parameters.

|  | Day1 | Day2 | Day3 |  | Day4 |  | Day5 |  |
| --- | --- | --- | --- | --- | --- | --- | --- | --- |
|  | Habituation | Fear learning | Contextual test | Cue test | 11840Hz test | 7000Hz test | White Noise test | Cue retest |
| Control | 10min | 3CS<br>CS: 20s<br>Without US | 3min | 2CS<br>CS:20s | 2CS<br>CS:20s | 2CS<br>CS:20s | 2CS<br>CS:20s | 2CS<br>CS:20s |
| Standard | 10min | 3CS-US<br>CS: 20s<br>US:0.7mA, 1s | 3min | 2CS<br>CS:20s | 2CS<br>CS:20s | 2CS<br>CS:20s | 2CS<br>CS:20s | 2CS<br>CS:20s |
| wFC | 10min | 3CS-US<br>CS: 20s<br>**US:0.5mA, 0.5s** | 3min | 2CS<br>CS:20s | 2CS<br>CS:20s | 2CS<br>CS:20s | 2CS<br>CS:20s | 2CS<br>CS:20s |
| sFC | 10min | 3CS-US<br>CS: 20s<br>**US:0.9mA, 2s** | 3min | 2CS<br>CS:20s | 2CS<br>CS:20s | 2CS<br>CS:20s | 2CS<br>CS:20s | 2CS<br>CS:20s |
| SL | 10min | 3CS-US<br>**CS: 5s**<br>US:0.7mA, 1s | 3min | 2CS<br>CS:20s | 2CS<br>CS:20s | 2CS<br>CS:20s | 2CS<br>CS:20s | 2CS<br>CS:20s |
| LL | 10min | 3CS-US<br>**CS: 80s**<br>US:0.7mA, 1s | 3min | 2CS<br>CS:20s | 2CS<br>CS:20s | 2CS<br>CS:20s | 2CS<br>CS:20s | 2CS<br>CS:20s |
| ST | 10min | 3CS-US<br>CS: 20s<br>US:0.7mA, 1s | 3min | 2CS<br>**CS:5s** | 2CS<br>**CS:5s** | 2CS<br>**CS:5s** | 2CS<br>**CS:5s** | 2CS<br>**CS:5s** |
| LT | 10min | 3CS-US<br>CS: 20s<br>US:0.7mA, 1s | 3min | 2CS<br>**CS:80s** | 2CS<br>**CS:80s** | 2CS<br>**CS:80s** | 2CS<br>**CS:80s** | 2CS<br>**CS:80s** |

wFC: weak fear conditioning group; sFC: strong fear conditioning group; SL: short learning group; LL: long learning group; ST: short test group; LT: long test group

### Measurement of freezing behavior

The total amount of freezing as a percentage of the total exposure time was used as a traditional measure of fear. Freezing was defined as a cessation of movement, with the exception of respiration, lasting for more than 1 s. The duration of freezing was then transformed into a percentage score representing the entire duration of the trial. Throughout all sessions, video recordings were used to evaluate freezing behavior with the *FreezeFrame* software (Actimetrics, Wilmette, IL). The software operates by assessing shifts in pixel luminance intensity across consecutive video frames (captured at a rate of 1–15 frames per second), and subsequently computing the changes in motion, as these two parameters show a linear correlation. A predefined threshold is applied to the data, thereby generating a percentage freezing score. In our experiment, during to an operating error, the frames were captured at a rate of 3.75 frames per second in the standard, control, SL, LL, ST, and LT groups, while at a rate of 15 frames per second in the sFC and wFC groups. Upon the conclusion of the experiments, we batch exported data and conducted analyses, with results directly transferred to an open *Excel* spreadsheet for 5 s intervals.

### Collection and identification of 22kHz vocalizations

USV data were collected using two M500-384 USB Ultrasound Microphones (Pettersson Elektronik AB, frequency range 0-150kHz) positioned directly above the floor in each chamber. These microphones were connected to computers via USB 2.0, and recordings were captured with Audacity 2.3 at a sampling rate of 384,000Hz in a 12-bit format.

For the detection and analysis of all USV recordings, we utilized the *DeepSqueak* toolbox based on MATLAB, a sophisticated artificial intelligence tool grounded in a cutting-edge regional convolutional neural network architecture (Faster-RCNN),which has demonstrated superior recognition rates compared to traditional detection methods [31]. Post-automated identification by the software, we performed manual screening via spectrogram analysis conducted by *DeepSqueak*, ensuring all USVs were accurately identified. The 22kHz USVs were evaluated based on specific criteria: we excluded items that did not fall within the 18-35kHz frequency range, displayed a waveform that was a single straight line or a diffuse point, or demonstrated unchanging intensity in energy. Seven trained researchers, blinded to the study’s purpose, manually reviewed the machine-recognized 22kHz USVs, strictly adhering to the above judgment criteria. The final identified 22kHz USV results were exported as .xlsx files, containing data on the start time, duration, and end time of each vocalization segment. A custom tool, developed in Python 3.6, was utilized to calculate the percentage of 22kHz calls per rat every 5 s throughout the experiment using these time data.

### Statistical analysis

All statistical analyses and graphical representations were conducted using GraphPad Prism 9. Prior to analysis, all data were examined for normal distribution and subjected to a Hartley’s test to assess the homogeneity of variance. No significant deviations were identified. Differences in the percentage of rats that emitted 22kHz USVs among groups were examined using the Mann-Whitney U test. A Student’s unpaired t-test was employed to compare two independent groups, and for multiple comparisons, we used a two-way repeated measures ANOVA with Tukey and Sidak’s *post hoc* tests. All data are presented as means ± standard errors of the means (SEMs). Differences were considered statistically significant at *p* < 0.05.

For percentage scores of freezing behavior at every 5 s interval and 22kHz vocalizations in each group, we conducted z-standardization separately for these two datasets and used the Pearson correlation to calculate the correlation between freezing behavior and 22kHz USVs within the same time period. To further depict the trends, we calculated the slope (*k*) of the data at corresponding time points of the CS and interstimulus interval (ISI) to represent the rate of change.

## Results

Our comprehensive results, encompassing the expression of 22kHz USV and freezing across eight distinct experimental conditions (control, standard, weak fear conditioning – wFC, strong fear conditioning – sFC, short CS fear learning – SL, long CS fear learning – LL, short CS cue test – ST, long CS cue test – LT), are illustrated via time series and cumulative data (see **Figures S1–S9** for details). We observed significant expressions of both 22kHz USV and freezing during fear learning and the cue test, with virtually no freezing or 22kHz USV evident in the contextual test. These findings suggest that all groups, excluding the control, successfully learned the association between CS and US. However, the degree of alignment between the two fear indicators was influenced by the intensity of the US and the duration of the CS during fear learning.

### 1. The percentage of rats emitting 22kHz calls is related to fear conditioning parameters

The percentage of rats emitting 22kHz calls during fear learning, cue tests, and fear generalization tests across all groups is tabulated in **Table 2**. It’s noteworthy that not all subjects emitted 22kHz calls, even after exposure to three footshocks during fear learning. However, we observed that the percentage of subjects emitting 22kHz calls, as well as the consistency of this bUSehavior, increased with the intensity of the footshocks. Among the four groups with mismatched CS durations between learning and tests (SL, LL, ST, LT), all except the LL group displayed lower percentages of rats emitting 22kHz calls in tests compared to the standard group. We employed Mann-Whitney U tests to evaluate the significance of the differences across groups (**Table 3**). As the intensity of the footshock escalated, the number of rats emitting 22kHz calls rose, although no significant difference was noted between the sFC group and the standard group. In groups with mismatched CS durations between learning and tests, we found fewer rats emitted 22kHz calls when the CS duration in the cue test exceeded that during fear learning.

**Table 2.**
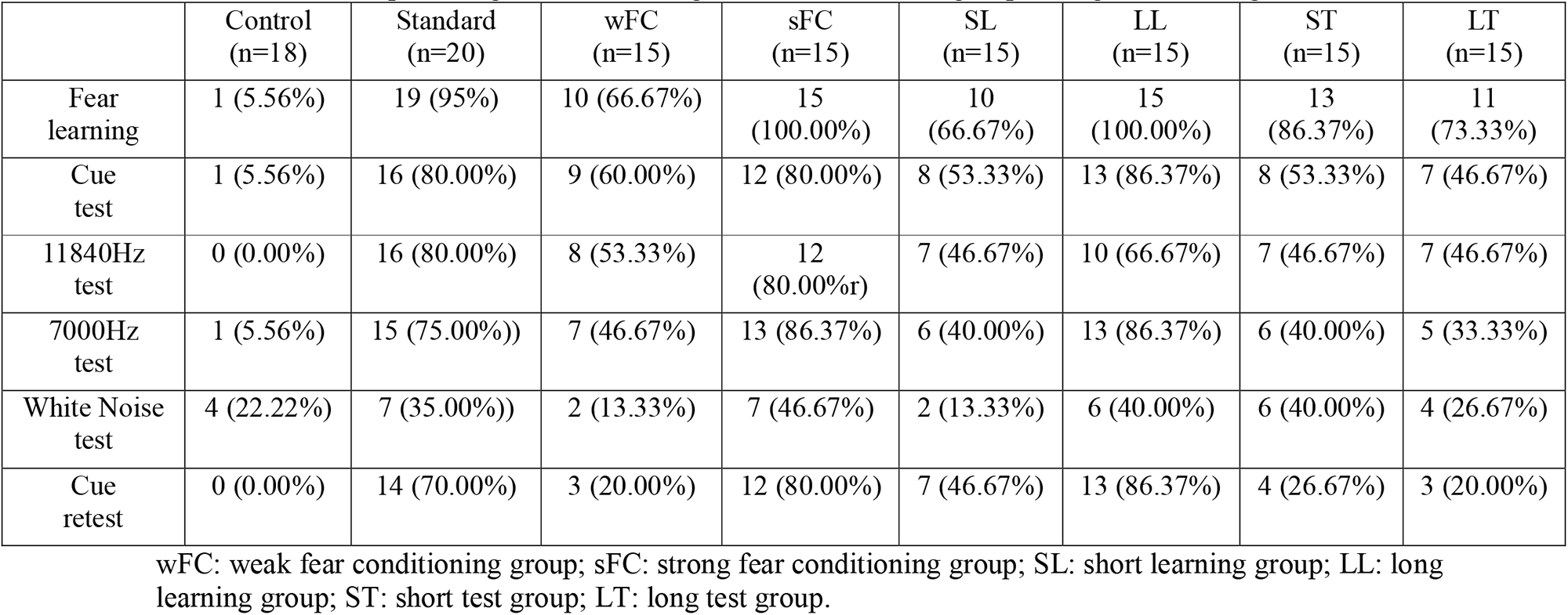
The percentage of rats emitting 22kHz calls in each group during fear learning and tests.

**Table 3.** Mann-Whitney U test’s results during the fear conditioning and fear generalization.

| A | B | A median | B median | Mann- Whitney U value | P value |
| --- | --- | --- | --- | --- | --- |
| Control vs | standard | 5.56 | 77.5 | 0 | 0.002 |
| Control vs | wFC | 5.56 | 50 | 2 | 0.007 |
| Standard vs | wFC | 77.5 | 50 | 4 | 0.024 |
| Standard vs | sFC | 77.5 | 80 | 12 | 0.39 |
| Standard vs | SL | 77.5 | 46.67 | 5 | 0.04 |
| Standard vs | LL | 77.5 | 86.37 | 13 | 0.45 |
| Standard vs | ST | 77.5 | 43.34 | 9 | 0.16 |
| Standard vs | LT | 77.5 | 40 | 4 | 0.02 |
| SL vs | LL | 46.67 | 86.37 | 5 | 0.03 |
| ST vs | LT | 43.34 | 40 | 13.5 | 0.50 |

### 2. US intensity influences the expression and matching of freezing and 22kHz USVs in fear conditioning

To elucidate the effects of US intensity on fear response, we compared the expression of freezing and 22kHz USVs between the control and standard groups. During the fear learning phase, distinct behavioral patterns were observed between the two groups (**Figure 2A**). The standard group exhibited a consistently increasing trend of freezing upon footshock administration, whereas the control group showed an immediate rise in freezing upon hearing the CS, followed by a steady decline^1^. Yet, no significant differences were found in total freezing between the two groups (Repeated measures ANOVA, *F* _(1, 36)_ = 0.98, *p* > 0.05, **Figure 2B**). During the cue test, although the standard group initially exhibited higher freezing than the control group (**Figure 2E**), this difference was not significant (Repeated measures ANOVA, *F* _(1, 36)_ = 1.71, *p* > 0.05; **Figure2F**). Nonetheless, a large number of 22kHz calls were observed exclusively in the standard group during both fear learning and the cue test. In fear learning, the standard group consistently emitted 22kHz calls after footshock, and from the second trial onwards, the rats began to emit 22kHz calls upon CS commencement (**Figure 2C**). During the cue test, the standard group emitted 22kHz calls after the end of the CS, which the control group did not (Figure 2G). Thus, significant differences in 22kHz USVs were observed between the control and standard groups during fear learning (Repeated measures ANOVA: *F* _(1, 36)_ = 45.29, *p* < 0.0001; **Figure 2D**) and the cue test (Repeated measures ANOVA: *F* _(1, 36)_ = 23.21, *p* < 0.0001; **Figure 2H**). Additionally, we found that for each group, the temporal correlation between 22kHz calling and freezing was related to US intensity. Higher US intensity led to more consistent trends in 22kHz USV and freezing during fear learning and the cue test (for detailed results see **Table 4–5**).

**Figure 2.**
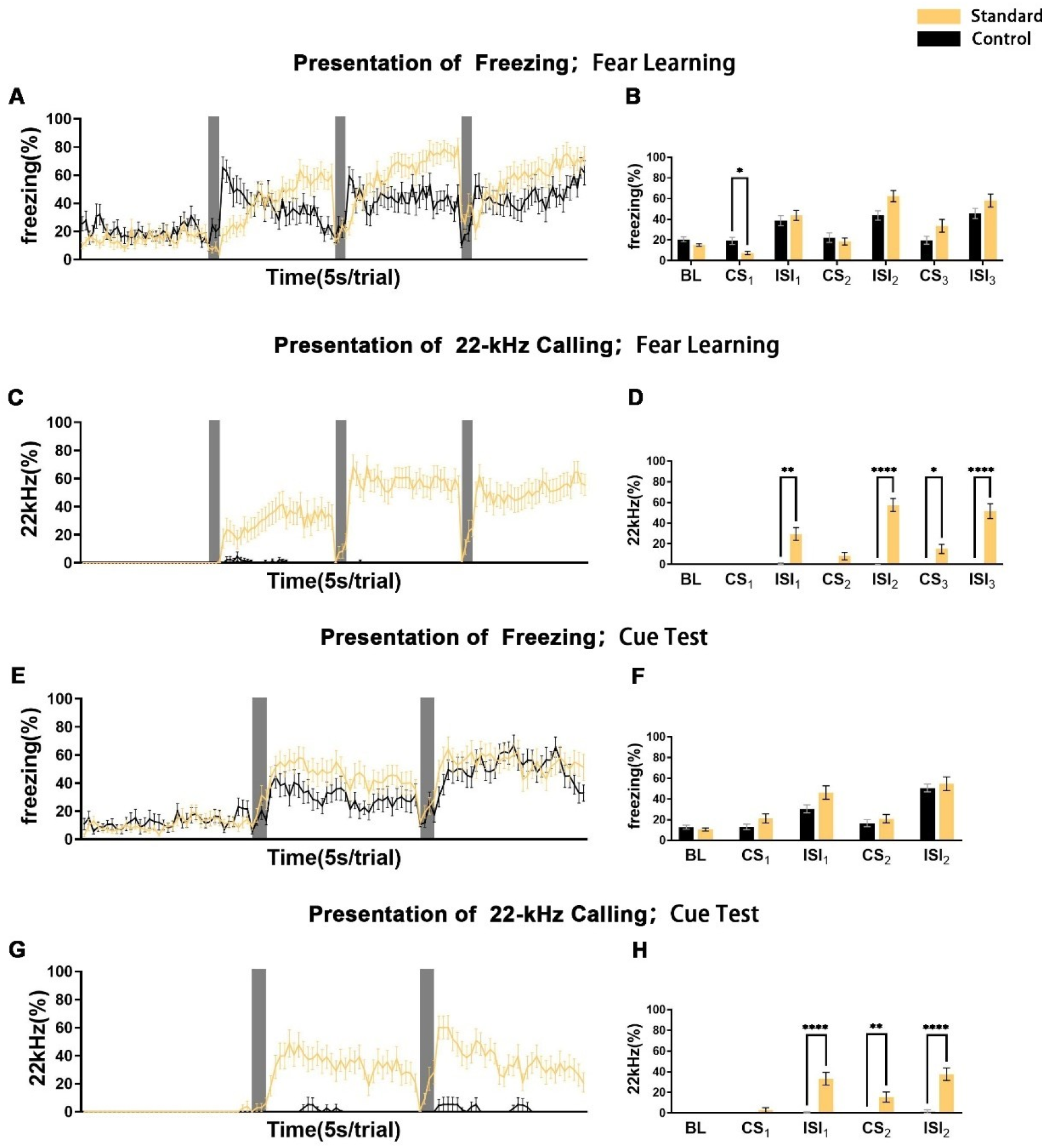
Comparison of 22kHz vocalizations and Body Freezing in Standard and Control Groups during Fear Learning and Cue Test. The graphs illustrate the manifestation of freezing (A, B), and 22kHz USVs (C, D) during fear learning, as well as the manifestation of freezing (E, F) and 22kHz USVs (G, H) during the cue test between the standard and control groups. In the left column (A, C, E, G), gray segments denote the CS presentation periods, and each point represents a mean score for a 5-second interval with its corresponding SEM. The right column (B, D, F, H) displays the cumulative values for baseline (BL), CS presentation (CS), and interstimulus interval (ISI) for each experiment. The standard group comprises 20 subjects and the control group comprises 18 subjects. Significance levels are indicated as follows: \**p*<0.05, \*\**p*<0.01, \*\*\**p*<0.001, \*\*\*\**p*<0.0001.

**Figure 3.**
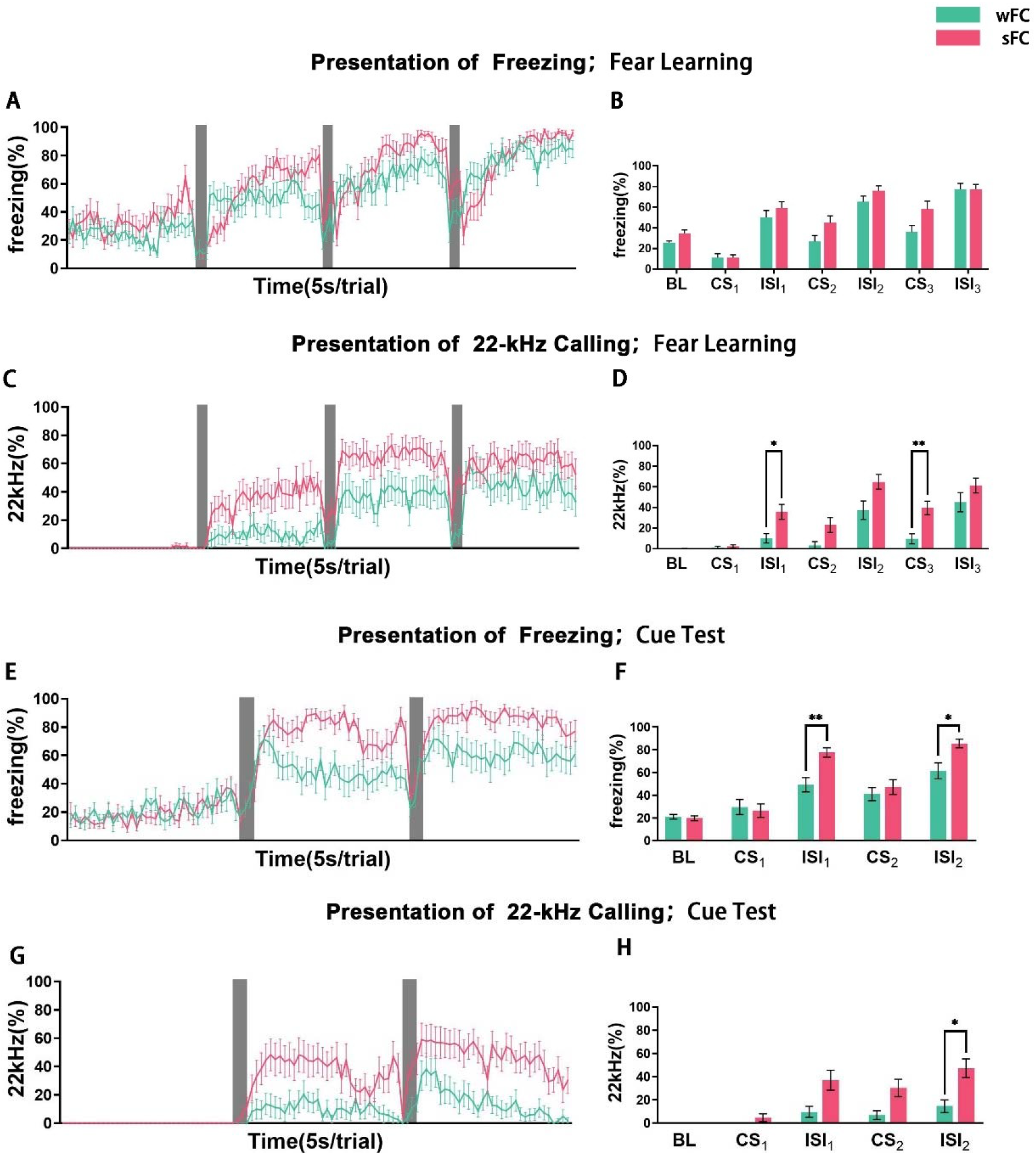
Comparison of 22kHz vocalizations and Body Freezing in Strong (sFC) and Weak (wFC) Fear Conditioning Groups during Fear Learning and Cue Test. The graphs present the expression of freezing (A, B), and 22kHz USVs (C, D) during fear learning, as well as the expression of freezing (E, F) and 22kHz USVs (G, H) during the cue test between the sFC and wFC groups. In the left column (A, C, E, G), gray segments represent the periods of CS presentation, and each point indicates a mean score for a 5-second interval with its corresponding SEM. The right column (B, D, F, H) showcases the cumulative values for baseline (BL), CS presentation (CS), and interstimulus interval (ISI) for each experiment. The sFC group and the wFC group each comprise 15 subjects. Significance levels are denoted as follows: \**p*<0.05, \*\**p*<0.01

**Table 4.** Pearson correlation coefficients between freezing and 22kHz calls (fear learning)

|  | Standard | sFC | wFC | LL | SL | LT | ST |
| --- | --- | --- | --- | --- | --- | --- | --- |
| Total | 0.880<sup>***</sup> | 0.794<sup>***</sup> | 0.865<sup>***</sup> | 0.905<sup>***</sup> | 0.746<sup>***</sup> | 0.807<sup>***</sup> | 0.855<sup>***</sup> |
| CS<sub>1</sub> | -0.907 | 0.767 | 0.189 | -0.43 | - | 0 | -0.04 |
| ISI<sub>1</sub> | 0.746<sup>**</sup> | 0.759<sup>**</sup> | -0.242 | 0.016 | 0.323 | 0.121 | -0.506 |
| ISI<sub>1</sub><br>(first 4 trials) | 0.076 | 0.922<sup>*</sup> | -0.791 | -0.399 | -0.341 | 0 | -0.544 |
| CS<sub>2</sub> | 0.539 | 0.948<sup>*</sup> | 0.226 | 0.925<sup>***</sup> | - | 0.712 | 0.79 |
| ISI<sub>2</sub> | -0.068 | 0.253 | 0.458 | -0.333 | 0.556<sup>***</sup> | 0.121 | -0.506 |
| ISI<sub>2</sub><br>(first 4 trials) | 0.230 | 0.875 | 0.866 | -0.356 | 0.342 | -0.044 | 0.305 |
| CS<sub>3</sub> | 0.621 | 0.967<sup>*</sup> | 0.591 | 0.937<sup>***</sup> | - | 0.842<sup>*</sup> | 0.817<sup>*</sup> |
| ISI<sub>3</sub> | 0.297 | 0.436 | -0.493 | -0.023 | 0.251 | -0.454 | -0.433 |
| ISI<sub>3</sub><br>(first 4 trials) | -0.356 | 0.886<sup>*</sup> | 0.383 | 0.732 | 0.938<sup>*</sup> | 0.553 | -0.302 |

\* $p < .05$ , \*\* $p < .01$ , \*\*\* $p < .001$

### 3. The duration-matching of CS between fear learning and tests influences the expression and matching of freezing and 22kHz USVs in fear conditioning

We first compared the SL and LL groups, which received shorter or longer CS durations respectively than the standard group during fear learning, to investigate the impact of varying CS on freezing and 22kHz USV. Both groups exhibited freezing and 22kHz calls during fear learning. Interestingly, in the LL group with longer CS duration, rats quickly froze and emitted 22kHz calls upon CS initiation in the second and third trials. Moreover, the LL group displayed a higher correlation between freezing and 22kHz USV than the SL group (Pearson correlation: LL: 0.91, SL: 0.74; **Figure 4A-B**). Despite the same footshock intensity, the variation in CS duration during fear learning led to a mismatch between freezing and 22kHz USV during the cue test, particularly in the first trial (**Figure 4C-D**). During the first CS, both the SL and LL groups froze at the beginning of CS playback (SL group’s *k*: 2.77; LL group’s *k*: 11.28) but did not emit 22kHz calls. Yet, after the second CS, both groups exhibited an increase in freezing and 22kHz USV at the onset of CS playback (SL group’s *k* of freezing: 5.11, *k* of 22kHz USV: 4.00; LL group’s *k* of freezing: 9.75, *k* of 22kHz USV: 7.94), and the trends became similar (Pearson correlation: LL: 0.98, SL: 0.85). No significant differences in freezing were observed among the SL, LL, and standard groups in the cue test (Repeated measures ANOVA: *F* _(2, 47)_ = 0.68, *p* > 0.05; **Figure 4E-F**), but significant differences were detected in 22kHz USV, particularly during ISI1 (Repeated measures ANOVA: *F* _(2, 47)_ = 4.73, *p* < 0.05; Tukey’s *post hoc* test: ISI_1_: standard and SL: Mean diff: 24.80, *p* < 0.01; standard and LL: Mean diff: 22.19, *p* < 0.05; ISI_2_: standard and SL: Mean diff: 21.25, *p* < 0.05; **Figure 4G-H**).

**Figure 4.**
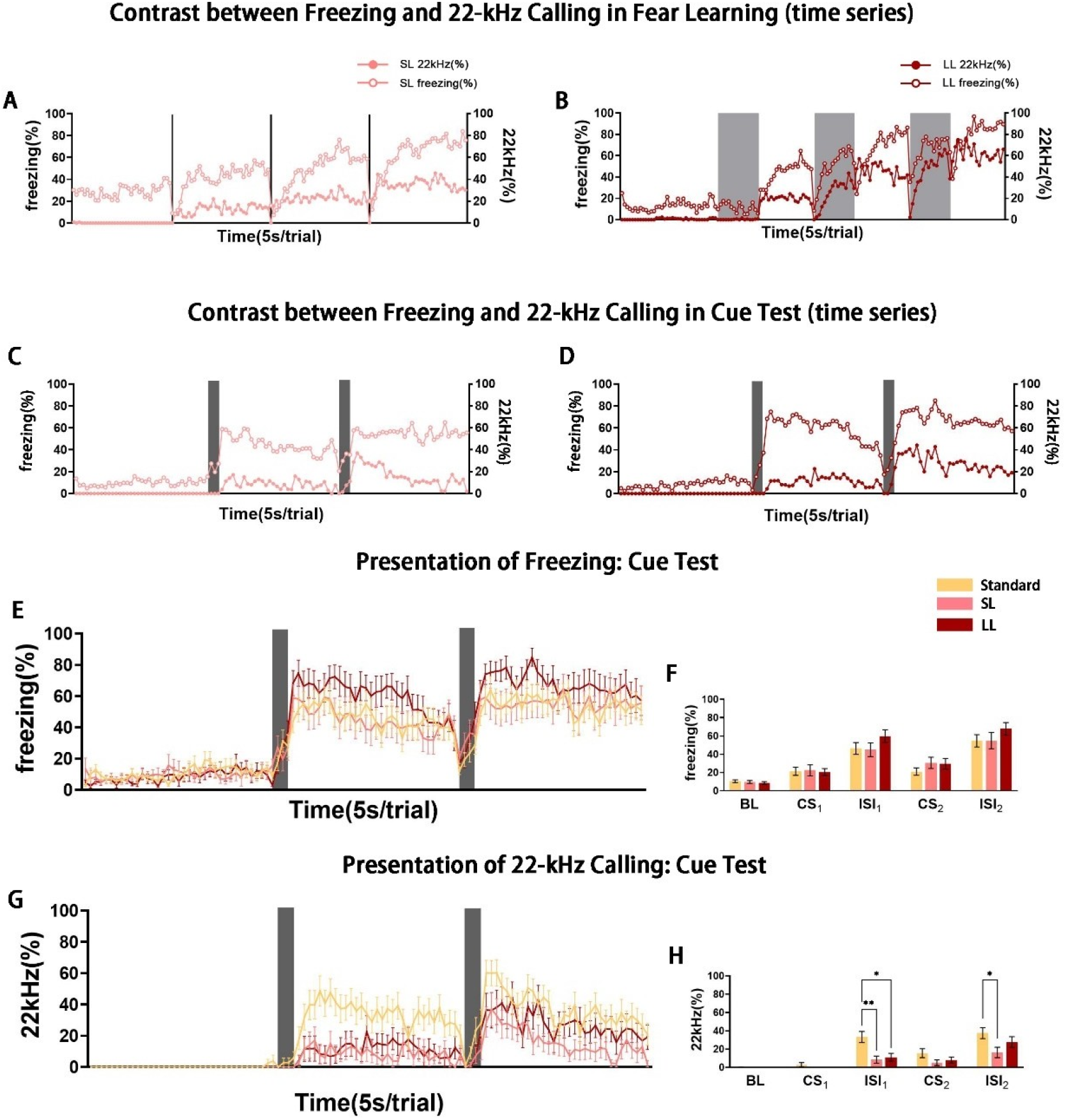
Comparison of 22kHz vocalizations and Body Freezing in Groups with Different CS Durations during Fear Learning and Cue Test, Compared to the Standard Group. Gray sections denote the CS presentation periods, and each point represents an average score over a 5-second interval, along with its corresponding SEM. The graphs showcase the expression of freezing and 22kHz USVs during fear learning (A, B), and the expression of freezing and 22kHz USVs (C, D) during the cue test for the short learning (SL) and long learning (LL) groups. Scores of freezing for every 5-second interval (left Y-axis) and the percentage of 22kHz vocalizations (right Y-axis) are compared and reported correspondingly. Figures E-H display the expression of freezing (E, F) and 22kHz USVs (G, H) during the cue test for the standard, LL, and SL groups. The right column (F, H) illustrates the cumulative values for baseline (BL), CS presentation (CS), and interstimulus interval (ISI) for each experiment. The SL group and the LL group each consist of 15 subjects, while the standard group includes 20 subjects. Significance levels are denoted as follows: \**p*<0.05, \*\**p*<0.01.

Next, to investigate the impact of varying CS duration on freezing and 22kHz USV during the cue tests, we compared the responses of the ST and LT groups. These groups had the same CS duration as the standard group during fear learning but had a shorter or longer CS duration than the standard group during the cue and generalization tests. For both freezing and 22kHz USV, the three groups exhibited similar trends during fear learning (Pearson correlation: standard: 0.88, LT: 0.80, ST: 0.85, *p* < 0.001). However, the ST group displayed more freezing (Repeated measures ANOVA: *F* _(2, 47)_ = 5.65, *p* < 0.01; Tukey’s *post hoc* test: ISI_3_: Standard and ST: Mean diff: 22.84, *p* < 0.05; **Figures 5A-B**) than the other two groups, while no significant differences in 22kHz USV were observed among the groups (Repeated measures ANOVA: *F* _(2, 47)_ = 0.94, *p* > 0.05; **Figures 5C-D**). In the cue test, we observed a discordance between freezing and 22kHz USV caused by the mismatch of CS duration. In the ST group, rats immediately froze after the conclusion of the first CS (CS1-ISI1 *k*: 45.61), maintaining a fluctuating increase at the initial four trials of ISI1 *k*: 0.97). However, the 22kHz calls did not immediately respond after the end of the first CS but continued to rise over time (CS1-ISI1 *k*: 4.22, the first four trials *k*: 4.08). In the second trial, the responses of the ST group rats to freezing and 22kHz calls rapidly appeared after the end of CS (CS_2_-ISI_2_ freezing’s *k*: 33.85, CS_2_-ISI_2_ 22kHz USV’ *k*: 19.21; **Figure 5E**). For the LT group, freezing increased approximately 20 seconds after the first CS onset, but the 22kHz USV was delayed by about 10 seconds before it occurred. During CS2, rats froze and emitted 22kHz calls at the onset of CS (first four trials in CS2: freezing’s *k*: 7.70, 22kHz USV’ *k*: 3.90), and the trend of change was highly similar (Pearson correlation: total CS_2_: 0.89, first 4 trials in CS_2_: 0.93; **Figure 5F**).

**Figure 5.**
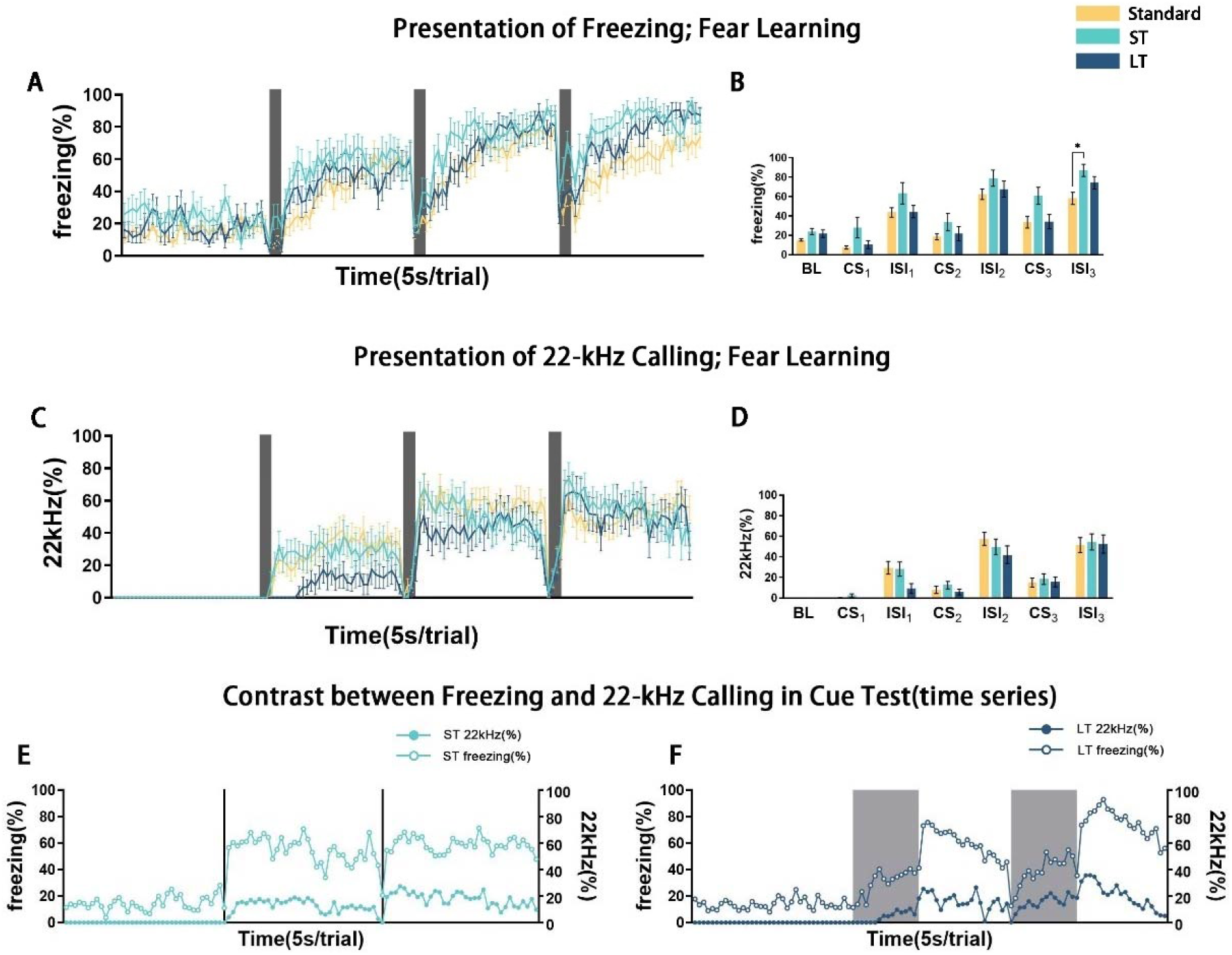
Comparison of 22kHz vocalizations and Body Freezing in Groups with Different CS Durations during the Cue Test, Compared to the Standard Group. The gray sections indicate the CS presentation periods, with each point representing an average score for a 5-second interval, along with its corresponding SEM. Figures A-D illustrate the expression of freezing (A, B) and 22kHz USVs (C, D) during the cue test for the standard, short test (ST), and long test (LT) groups. The right column (B, D) displays the cumulative values for baseline (BL), CS presentation (CS), and interstimulus interval (ISI) for each experience. The graphs in E and F represent the expression of freezing and 22kHz USVs during the cue test for the ST and LT groups. Scores of freezing for each 5-second interval (left Y-axis) and the percentage of 22kHz vocalizations (right Y-axis) are compared and reported accordingly. The ST group and LT group each consist of 15 subjects, while the standard group includes 20 subjects. Significance levels are indicated as follows: \**p*<0.05.

### 4. Differentiation of freezing and 22kHz USV in duration of cue was evident in generalization tests

Subsequent to the cue test, we conducted three rounds of fear generalization tests and a single round of cue retest for each group to explore the association between freezing and 22kHz USV in the context of fear generalization. Detailed time-series data for freezing and 22kHz USV during the generalization tests for all groups can be found in **Figures S6-9**. Notably, despite the high level of freezing exhibited by each group in the 11840Hz and 7000Hz generalization tests and the cue retest, the freezing response was not as pronounced during the White Noise test. Furthermore, the frequency of 22kHz USV diminished during the fear generalization tests. In the cue retest, the standard and sFC groups maintained strong expressions after the CS, whereas the 22kHz USV curve for the other groups remained relatively flat, exhibiting only a low peak.

Upon comparison across groups, we observed that groups subjected to different US intensities displayed identical levels of freezing during the generalization tests, however, 22kHz USV responses varied considerably. Specifically, no significant differences were found between the standard and control groups in all generalization tests and the cue retest. Nonetheless, a significant divergence in 22kHz USV between these two groups emerged in the 11840Hz and 7000Hz generalization tests, and in the cue retest (see **Table 6**). Moreover, the sFC group demonstrated higher freezing compared to the wFC group in the 11840Hz and 7000Hz generalization tests, with no difference in the white-noise test or the cue retest. Conversely, a significant disparity between the sFC and wFC groups was discernible in all generalization tests and the cue retest concerning 22kHz USV (see **Table 7**). Finally, irrespective of whether there were differences in CS duration during fear learning or during the cue test, no significant discrepancies in 22kHz USV between groups were noted in all generalization tests and the cue retest. However, differences in freezing were observed among the standard, ST, and LT groups during the white-noise test, with no significant differences noted in other groups (see **Tables 8–9**). The mismatch of CS duration did not seem to discriminate between freezing and 22kHz USV during fear generalization.

**Table 5.** Pearson correlation coefficients between freezing and 22kHz calls (cue test)

|  | Standard | sFC | wFC | LL | SL | LT | ST |
| --- | --- | --- | --- | --- | --- | --- | --- |
| Total | 0.925<sup>***</sup> | 0.950<sup>***</sup> | 0.704<sup>***</sup> | 0.835<sup>***</sup> | 0.735<sup>***</sup> | 0.871<sup>***</sup> | 0.891<sup>***</sup> |
| CS<sub>1</sub> | 0.757 | 0.904<sup>*</sup> | 0 | 0 | 0 | 0.683<sup>**</sup> | - |
| ISI<sub>1</sub> | 0.691 | 0.844<sup>***</sup> | 0.062 | 0.386 | 0.201 | 0.59 | 0.552 |
| ISI<sub>1</sub><br>(first 4 trials) | 0.948<sup>*</sup> | 0.990<sup>**</sup> | -0.641 | 0.175 | -0.801 | 0.787 | 0.468 |
| CS<sub>2</sub> | 0.972<sup>*</sup> | 0.909<sup>*</sup> | 0.967<sup>*</sup> | 0.980<sup>*</sup> | 0.853<sup>*</sup> | 0.897<sup>**</sup> | - |
| ISI<sub>2</sub> | 0.381 | 0.534<sup>**</sup> | 0.372 | 0.745<sup>***</sup> | -0.048 | 0.707 | 0.331 |
| ISI<sub>2</sub><br>(first 4 trials) | 0.037 | -0.265 | 0.566 | 0.573 | 0.693 | 0.647 | 0.418 |

\* $p < .05$ , \*\* $p < .01$ , \*\*\* $p < .001$

**Table 6.**
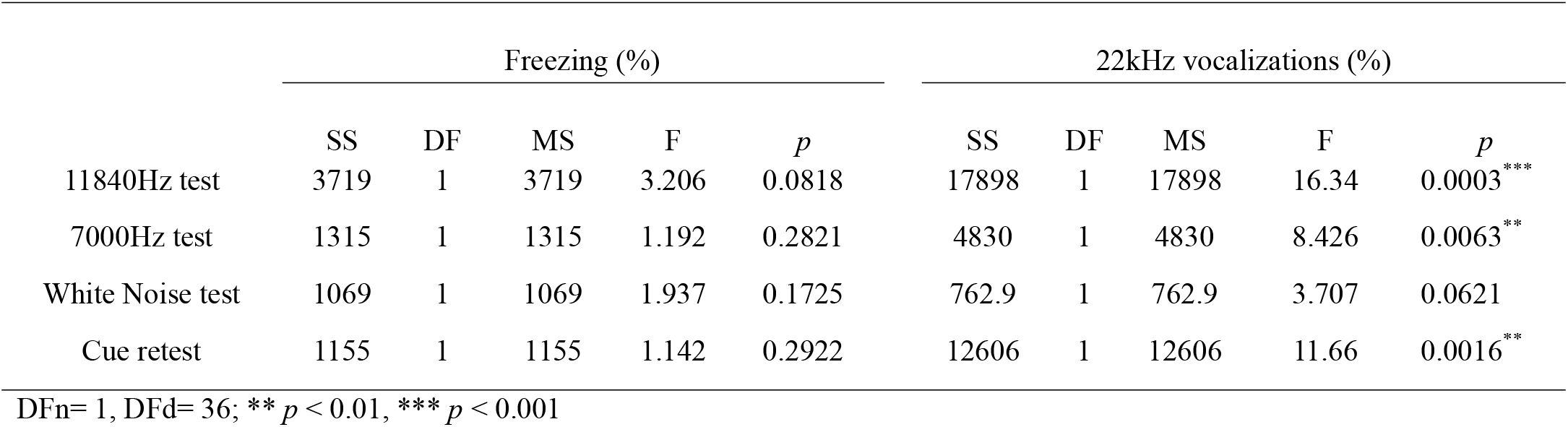
Repeated measures ANONA between standard and control groups during the generalization test and cue retest phase.

|  | Freezing (%) |  |  |  |  | 22kHz vocalizations (%) |  |  |  |  |
| --- | --- | --- | --- | --- | --- | --- | --- | --- | --- | --- |
|  | SS | DF | MS | F | *p* | SS | DF | MS | F | *p* |
| 11840Hz test | 3719 | 1 | 3719 | 3.206 | 0.0818 | 17898 | 1 | 17898 | 16.34 | 0.0003*** |
| 7000Hz test | 1315 | 1 | 1315 | 1.192 | 0.2821 | 4830 | 1 | 4830 | 8.426 | 0.0063** |
| White Noise test | 1069 | 1 | 1069 | 1.937 | 0.1725 | 762.9 | 1 | 762.9 | 3.707 | 0.0621 |
| Cue retest | 1155 | 1 | 1155 | 1.142 | 0.2922 | 12606 | 1 | 12606 | 11.66 | 0.0016** |

DFn= 1, DFd= 36; \*\* $p < 0.01$ , \*\*\* $p < 0.001$

**Table 7.** Repeated measures ANONA between wFC and sFC groups during the generalization test and cue retest phase.

|  | Freezing (%) |  |  |  |  | 22kHz vocalizations (%) |  |  |  |  |
| --- | --- | --- | --- | --- | --- | --- | --- | --- | --- | --- |
|  | SS | DF | MS | F | *p* | SS | DF | MS | F | *p* |
| 11840Hz test | 10934 | 1 | 10934 | 8.913 | 0.0058** | 19957 | 1 | 19957 | 15.37 | 0.0005*** |
| 7000Hz test | 15259 | 1 | 15259 | 13.88 | 0.0009*** | 13371 | 1 | 13371 | 14.89 | 0.0006*** |
| White Noise test | 1297 | 1 | 1287 | 1.962 | 0.1723 | 1412 | 1 | 1412 | 6.994 | 0.0133* |
| Cue retest | 1873 | 1 | 1873 | 1.454 | 0.2380 | 19605 | 1 | 19605 | 17.40 | 0.0003*** |

DFn= 1, DFd= 28; \* $p < 0.05$ , \*\* $p < 0.01$ , \*\*\* $p < 0.001$

**Table 8.**
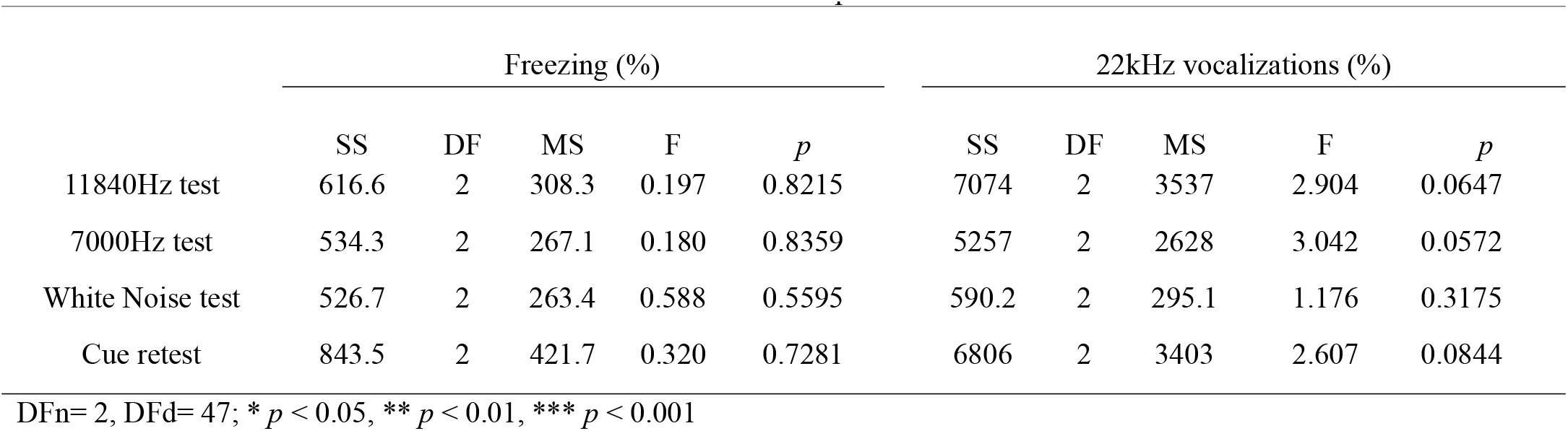
Repeated measures ANONA among groups with different CS durations in fear learning during the generalization test and cue retest phase.

**Table 9.** Repeated measures ANONA among groups with different CS durations in cue test during the generalization test and cue retest phase.

|  | Freezing (%) |  |  |  |  | 22kHz vocalizations (%) |  |  |  |  |
| --- | --- | --- | --- | --- | --- | --- | --- | --- | --- | --- |
|  | SS | DF | MS | F | *p* | SS | DF | MS | F | *p* |
| 11840Hz test | 90.23 | 2 | 45.11 | 0.030 | 0.9704 | 6967 | 2 | 3483 | 2.355 | 0.1060 |
| 7000Hz test | 2823 | 2 | 1411 | 0.954 | 0.3927 | 269.9 | 2 | 135 | 0.1053 | 0.9003 |
| White Noise test | 10340 | 2 | 5170 | 6.396 | 0.0035** | 114.8 | 2 | 57.41 | 0.1868 | 0.8302 |
| Cue retest | 4328 | 2 | 2164 | 1.479 | 0.2382 | 5501 | 2 | 2751 | 1.716 | 0.1909 |

DFn=2, DFd= 47; \* $p < 0.05$ , \*\* $p < 0.01$ , \*\*\* $p < 0.001$

## Discussion

This study probes the interrelationship and potential characterization of freezing behavior and 22kHz USVs within the fear conditioning paradigm. It scrutinizes the temporal sequence of freezing and 22kHz USVs under diverse conditions in this paradigm, focusing on the impact of various parameters on fear expression in rodents, and the intricate interactions between freezing and 22kHz USVs under each condition.

Our results posit that both freezing and 22kHz USVs provide unique vantage points for representing fear. Notably, manipulation in the intensity of the US and incongruities in the duration of the CS between fear learning and the cue test instigate a divergence between these two measures. Interestingly, rats only produced 22kHz USVs when subjected to shocks, whereas CS stimulation alone could precipitate freezing behavior, particularly in rats with previous experiences of forced immobility, such as restraint. Upon manipulation of shock intensity during fear learning, cue test, and generalization tests, freezing behavior remained invariant, unable to discern the magnitude of the threat. Conversely, 22kHz USVs demonstrated a greater sensitivity to the variations. Moreover, after experiencing fear learning at different spatio-temporal distances, rats were able to identify cognitive mismatch of threat cues in their 22kHz USVs during cue tests. These vocalizations typically followed freezing but no differences in freezing behavior were discernable, as these responses were predominantly automatic and conservative. Collectively, these data suggest that 22kHz USVs might be indicative of a rat’s subjective fear experience, whereas freezing likely embodies an automatic defensive response, which is consistent with LeDoux’s “two-system” model of fear.

While the debate surrounding LeDoux’s “two-system” framework and the nature and definition of fear is primarily centered on human subjects, the importance of animal models in fear research and the implications of updating the concept of fear for animal experiments cannot be overlooked [32]. Recent studies have expanded from focusing on freezing to include more active defensive behaviors, such as flight or active avoidance, to discuss a range of response patterns in rats facing threats [33–37]. As a crucial part of “conscious fear,” the subjective fear in animals plays a vital role in our understanding of human fears. The content of 22kHz calling comprises emotional and referential or descriptive information about threat sources and their characteristics [38]. We propose that it is more appropriate to apply subjective emotion reports to the measurement of fear representation. Recent research found that the 22kHz USVs emitted during flight is shorter, louder, and exhibits a higher peak frequency than that emitted during freezing [39]. This implies a stronger USV emission accompanies an active fear response. Findings in social learning of fear also indicate that 22kHz USVs may represent subjective fear signals in fear conditioning, while freezing is a defensive response. For example, naive rats will learn the fear response to CS from rats previously shocked during CS presentation, which includes fright or freezing [40, 41]. However, this social transmission of fear has only been observed behaviorally – relevant studies on 22kHz calling have not found that naive rats can directly learn 22kHz USVs from shocked rats [42, 43]. Yet, rats with prior experience can be awakened by electric shock in the test [44]. This aligns with the description of the subjective cognitive loop in LeDoux’s “two-system” framework – the perception of fear requires the accumulation and integration of past experiences. Indeed, 22kHz USVs represent a broad range of emotional states in rats and are interpreted as different negative emotions in different contexts [45–47]. Despite that 22kHz calling observed in fear conditioning is already recognized as a fear indicator, some argue it represents anxiety [13, 48]. However, considering the substantial overlap between fear and anxiety at the neuroanatomical and pharmacological level [49, 50], using anxiolytic drugs to suppress 22kHz vocalizations in fear conditioning paradigms in rats does not entirely account for these vocalizations as anxiety representations. Additionally, anxiety is an emotion for coping with potential threats, in preparation for future uncertainty [51]. Our findings, however, show that in fear conditioning, the occurrence of 22kHz USVs is linked with actual threats (footshocks), and the frequency of these vocalizations becomes more pronounced with stronger shocks. Most importantly, 22kHz USVs typically appears in large numbers after the end of the CS during cued tests. This outcome suggests that 22kHz calling is a conditioned response in fear conditioning, representing a measure of fear rather than anxiety, much like freezing.

In our results, although both 22kHz USVs and freezing behavior reflect fear states in rats during fear conditioning, their manifestations differ. The emergence of 22kHz vocalizations during fear conditioning can also be modulated by varying contextual factors. Wöhr et al. found that the frequency of 22kHz USV escalates with increased US intensity during fear learning and significantly attenuates during testing [16], which aligns with our findings. They also found a divergence of freezing and 22kHz USVs in fear learning and the cue test with a 0.5 mA footshock. According to our results, although the total freezing levels were similar for the wFC and sFC groups, the sFC group displayed higher freezing levels during inter-stimulus intervals (ISI), whereas the wFC group consistently had fewer 22kHz USVs across all tests, indicating that local behavioral differences can be discerned when US intensities vary. In a study on early stress enhancing fear, the increased freezing response of the early stress group was only manifested in the cue test, while a large number of 22kHz USVs in the early stress group could be found in both the fear learning phase and the cue test [52]. Another study showed that freezing behavior might not always accompany 22kHz USVs, but when 22kHz USVs occur, they are usually accompanied by freezing behavior [53]. These evidences together further suggest that 22kHz USVs are more sensitive as a fear indicator.

Regarding the mismatch of CS duration between fear learning and cue testing, previous studies found that freezing isn’t significantly affected by duration variables and can still achieve levels similar to fear learning in the early stages of extinction [54]. We also found that the mismatch of CS duration had no effect on freezing, which seemed to suggest that rats would ignore the time attribute of the CS when they were afraid. On the other hand, our results show that the 22kHz USVs in the cue test of all groups that do not match fear learning and cue test in CS durations were less than that of the standard group. Especially noteworthy was the SL and ST groups, which deviated significantly from the standard group. Furthermore, in the first ISI, we observed a gap between the start times of freezing and 22kHz calling in all mismatch groups. However, in the second ISI, freezing and 22kHz calling occurred simultaneously. This suggests that rats can identify whether CS attributes are consistent with past experiences during testing and communicate this through 22kHz calling. Research involving goldfish and human subjects supports the notion that organisms prospectively estimate the CS duration during testing [55, 56]. This perspective has implications for extinction since employing a CS duration different from the acquisition makes extinction easier to implement [57]. It should be noted that freezing usually occurs as CS starts, while 22kHz USV usually occurs after freezing, which is similar to the process in which the defensive loop participates in promoting the cognitive loop to perceive fear in the “two-system” framework. In addition, different from the results of Fanselow et al.’s study [23], this study is more similar to the earlier structure of aversion and fear learning research, that is, the shorter the CS-US interval is, the less significant the learning effect is [49, 58]. The performance of the SL group in the cue test supports this view. The ST group also showed lower 22kHz USV in the cue test. A potential support comes from the evidence that the shorter CS duration in the fear extinction, the more reduction in fear expression [56, 59].

The separation of 22kHz USV and freezing is not only observed in the fear conditioning, but also exists in the fear generalization and cue retest. Fear generalization can not only measure the fear degree of rats, but also measure the discrimination of rats in similar cues with the final cue retest. The three rounds of generalized tones we selected had some correlation with 9kHz. First of all, 11840Hz and 7kHz of pure tone are respectively the first and the second overtone of 9kHz, so the tone attribute is symmetrical and harmonious. Secondly, the dominant frequency of the white-noise complex sound we used was near 9kHz, by which we hoped to test the rats’ generalization ability to the complex sound stimulus. The performance of 22kHz USV and freezing in pure tone of rats is similar to that of cue test, which is consistent with the conclusion of Aizenberg and Geffen’s study [60], that is, when the physical characteristics of the two kinds of sound stimuli are similar, the fear generalization of sound stimuli is more likely to occur. However, in the white-noise test, the fear reaction of each group was significantly reduced, whether it was freezing or 22kHz USV. This is different from the previous understanding of white noise in fear research [61, 62]. We propose that, on the one hand, white noise, as a compound sound, is not similar to pure tone features, and consideration of the stimulus similarity dimension of fear generalization will indeed lead to a weaker response. On the other hand, several rounds of tests have been conducted before the white noise test, which may improve the rats’ ability to discriminate threat cues. For the mismatch of CS group, the footshock they received was of the same specification as the standard group, so no difference was found between them and the standard group in the generalization test. This supports that fear generalization is mainly affected by threat intensity [63]. However, between groups with different US intensities, the higher the US intensity, the more stable and massive the 22kHz calls. Freezing tends to converge with generalization testing. This may be a kind of habituation, which eliminates the influence of CS on the fear response without affecting the expression of corresponding memory components [64]. However, in a human fear generalization study, it was found that such similar cues could not completely eliminate the fear of previous stimuli [65]. Therefore, this further proves that freezing is more likely to be an automated response, while 22kHz USV is more likely to reflect the subjective fear of rats.

The body of evidence presented herein substantiates that, within the fear response of rats, both freezing behavior and 22kHz vocalizations may be elicited by two separate systems. This contradicts the conventional understanding of a “single system” for fear, leaning more towards LeDoux’s “two-system” framework. In fact, when comparing with findings from human studies, we discern many functional similarities in rats’ USVs (analogous to human subjective reports) and freezing behavior (paralleling human physiological and defensive responses) [66–68].

In summary, by applying the “two-system” framework to rodent models, we propose that freezing behavior is likely an automated response, while 22kHz vocalizations serve as a subjective fear indicator in rats.

### Limitations of the study

While our study provides compelling insights into fear conditioning paradigms in rodents, it is not without its limitations. Firstly, extant literature has highlighted the presence of individual differences in behavioral phenotypes among rats participating in fear conditioning experiments [58, 69]. Although our study presents a variety of combinations of freezing and 22kHz calling, our main objective was to examine their performance and relationship within the fear conditioning context. Future work could delve deeper into the implications of these diverse phenotypes.

Secondly, while the frequency of 22kHz calling can provide an insight into the subjective fear experience of rats, a ’ceiling effect’ is apparent at high fear conditioning levels. Moreover, USVs are rich in information, encompassing various features such as intensity, amplitude, and complex tone. Future research could aim to decode USVs more comprehensively, further illuminating their significance.

Lastly, the distribution of the percentage data may not adhere to a normal distribution, potentially introducing some bias into the results of the repeated measure ANOVA. Despite this, we chose to use parametric tests for the analysis of USVs data, following the precedent set by previous studies.

Despite these limitations, our findings are promising and encourage a reconsideration of the roles and implications of freezing and USVs in rodent fear conditioning models, which contributes fresh evidence to the ongoing discussion around the definition of fear from the perspective of animal experiments.

## Data availability

All data generated to support the findings of this study are available at https://data.mendeley.com/datasets/w8hjfsn4bw.

## Author contributions

BY and BZ conceived of the present project, designed the study and collected the data. LB and DY assisted in analysing the data. BZ wrote the first draft of the manuscript. BY critically revised the manuscript. All authors approved the final version of the manuscript.

## Supplemental Figures

**Figure S1:**
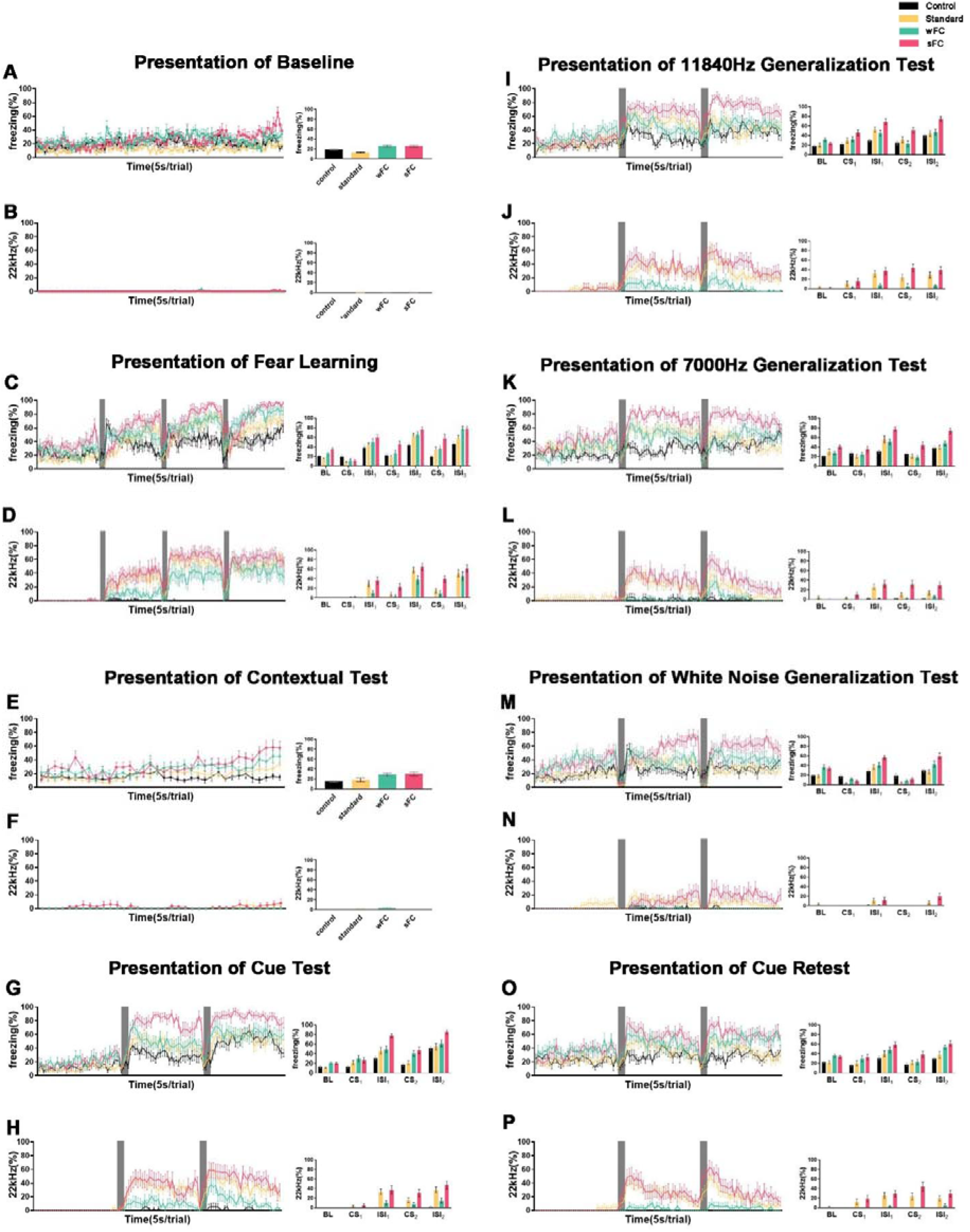
Variations in 22kHz vocalizations and Body Freezing with Different Intensity Levels of US, Compared with Control Conditions

**Figure S2:**
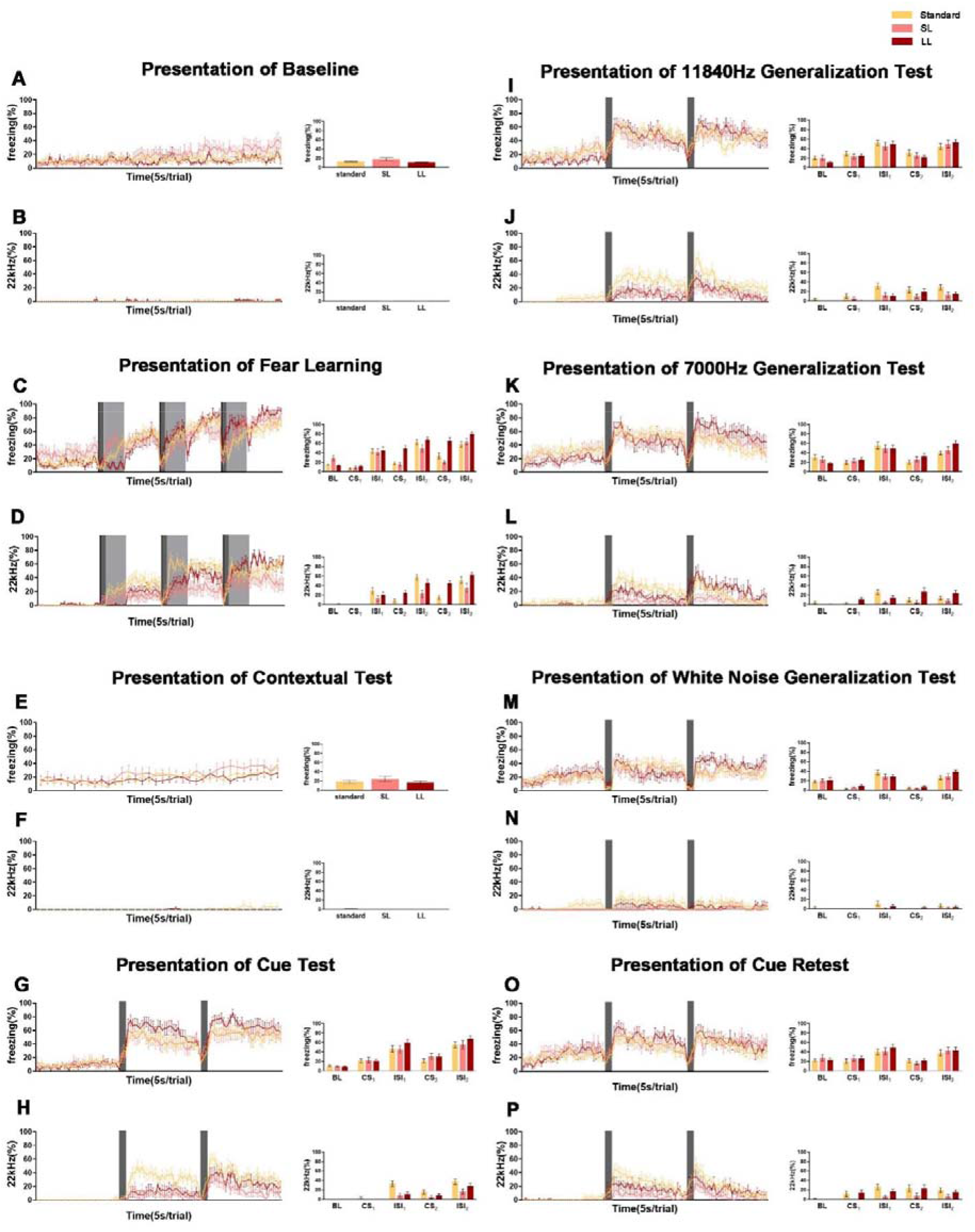
Variations in 22kHz vocalizations and Body Freezing with Different CS Durations during Fear Learning, Compared with Standard Conditions

**Figure S3:**
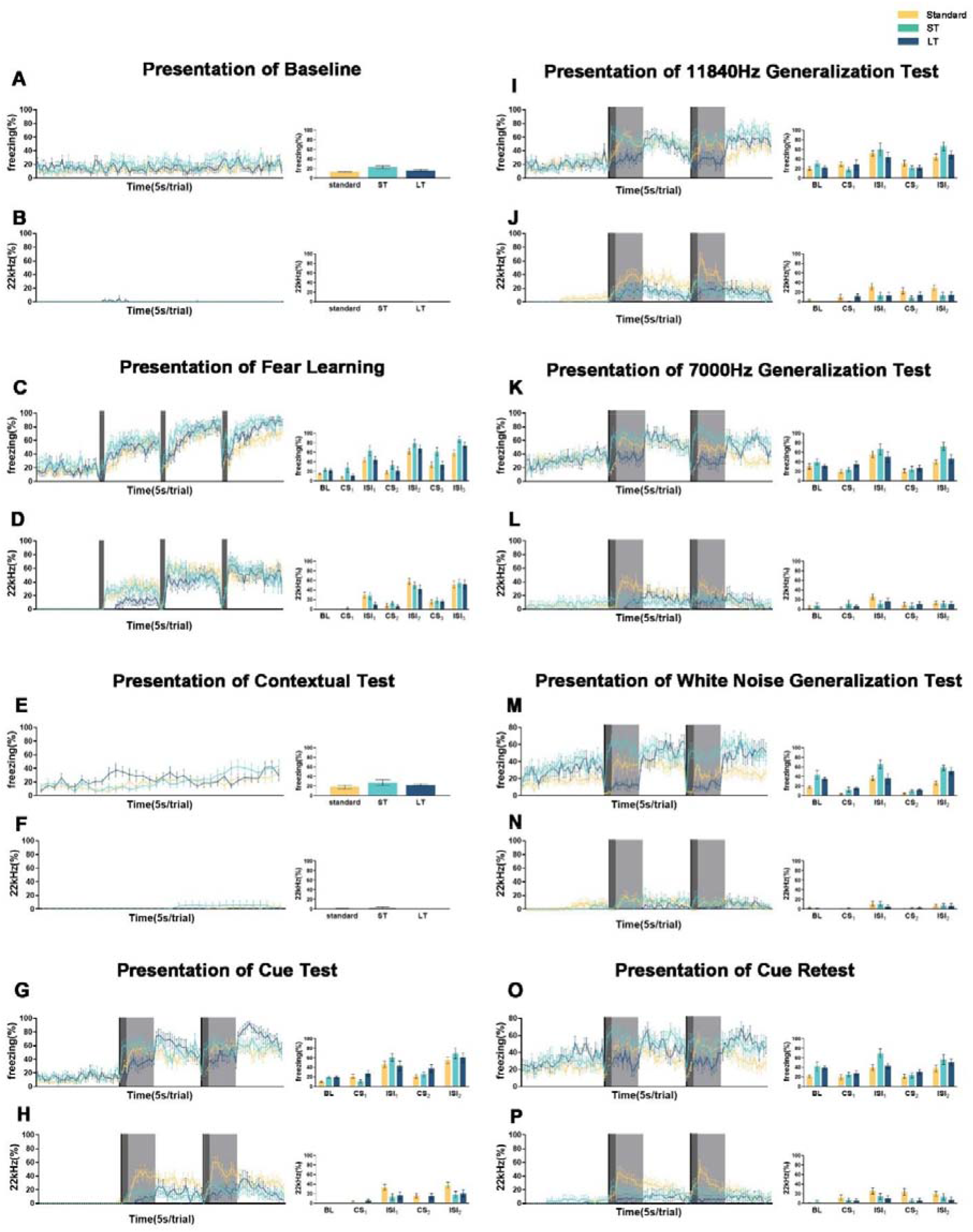
Variations in 22kHz vocalizations and Body Freezing with Different CS Durations during Cue Test, Compared with Standard Conditions

**Figure S4:**
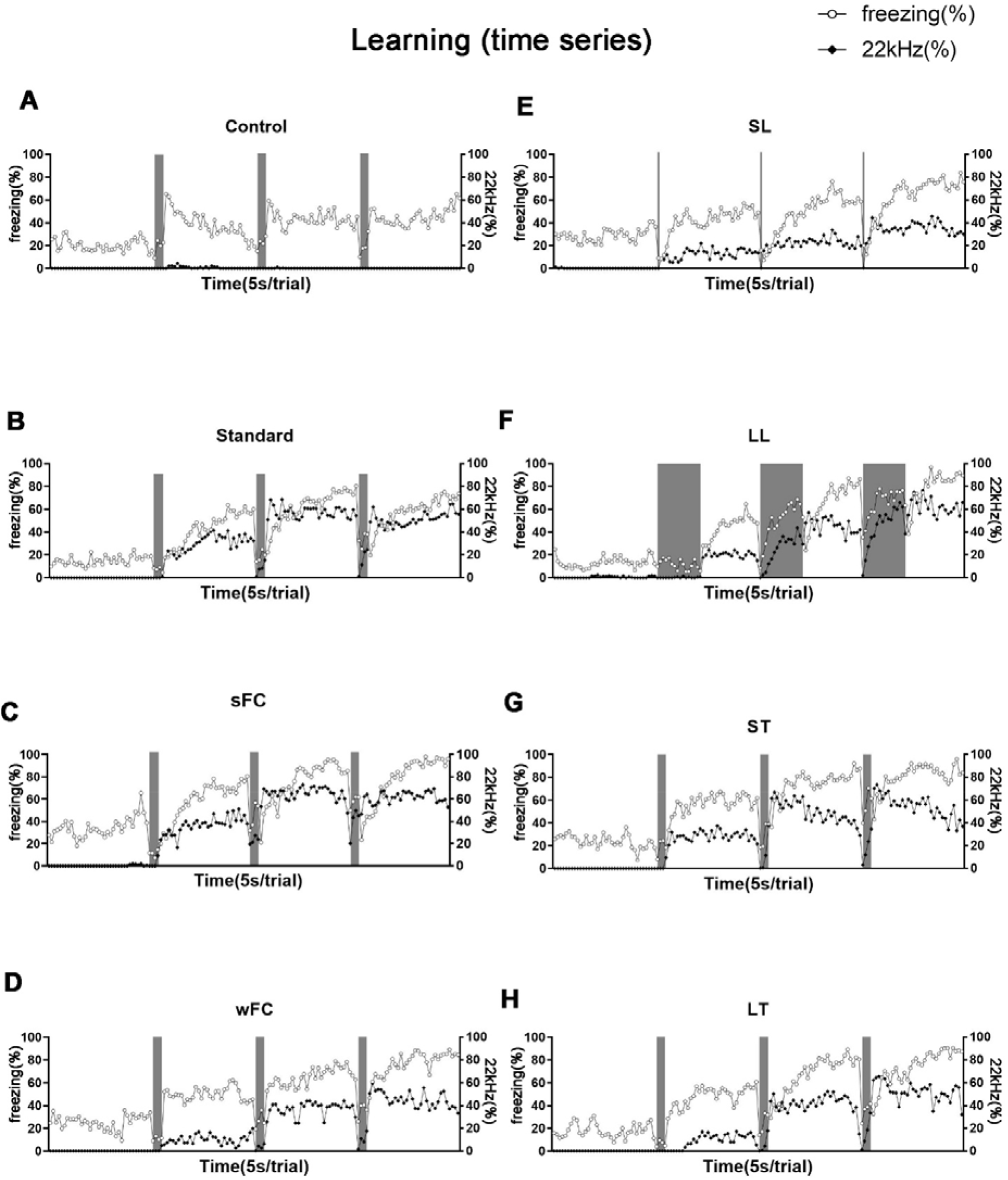
Temporal Analysis of 22kHz vocalizations and Body Freezing during Fear Learning Across Diverse Experimental Conditions, Including the Control Group

**Figure S5:**
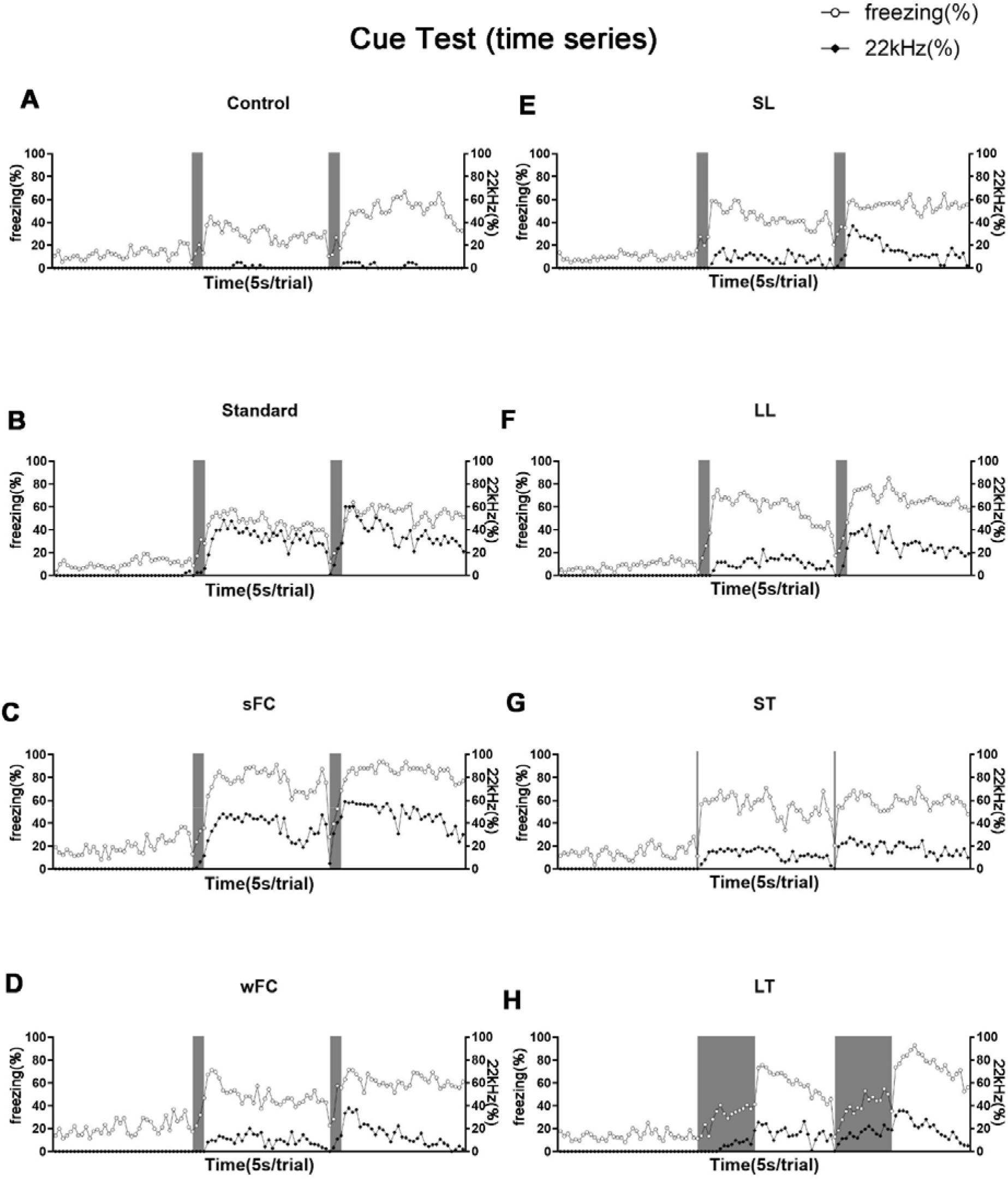
Temporal Analysis of 22kHz vocalizations and Body Freezing during Cue Test across Diverse Experimental Conditions, Including the Control Group

**Figure S6:**
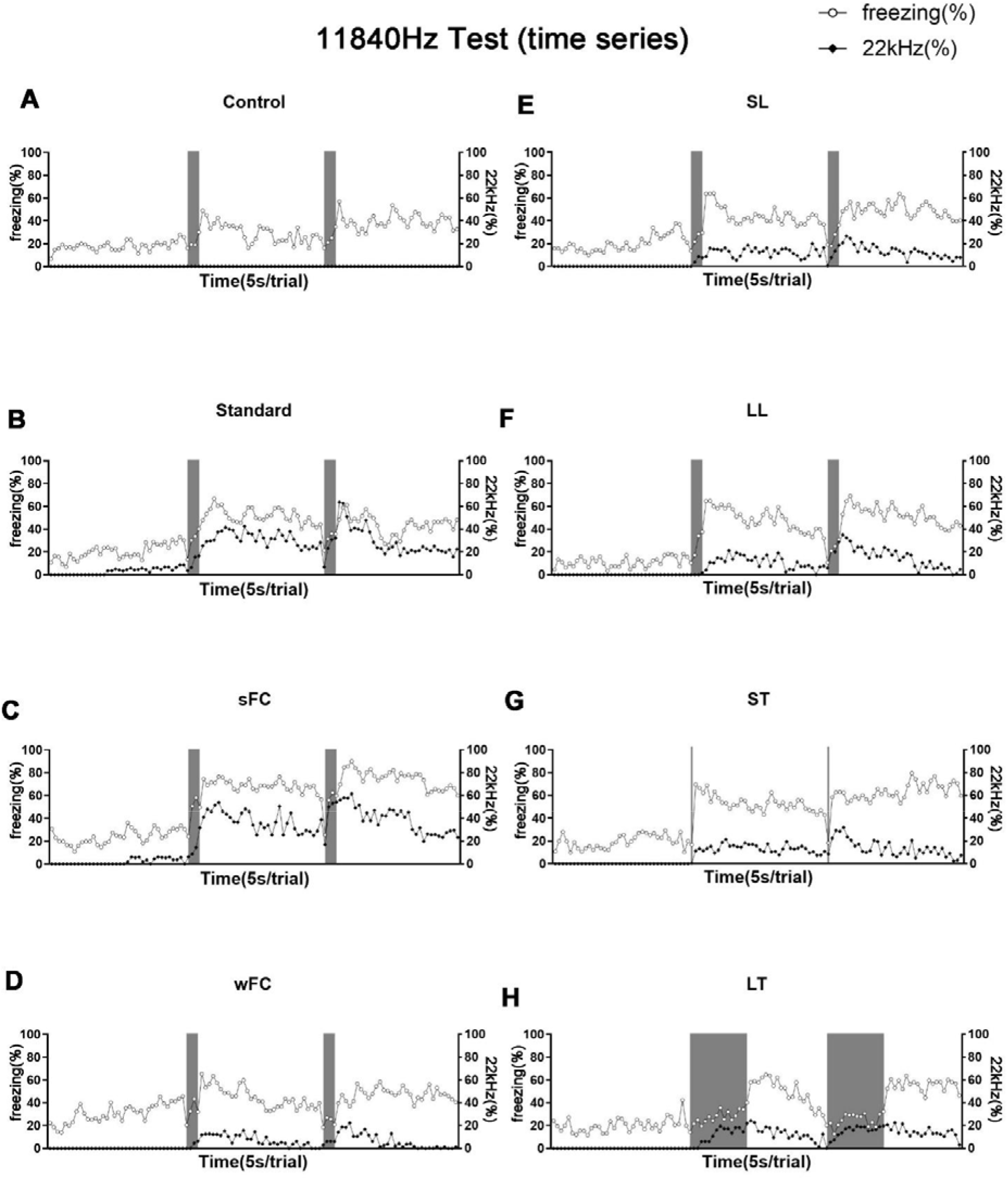
Temporal Analysis of 22kHz vocalizations and Body Freezing during 11840Hz Cue Generalization Test Across Diverse Experimental Conditions, Including the Control Group

**Figure S7:**
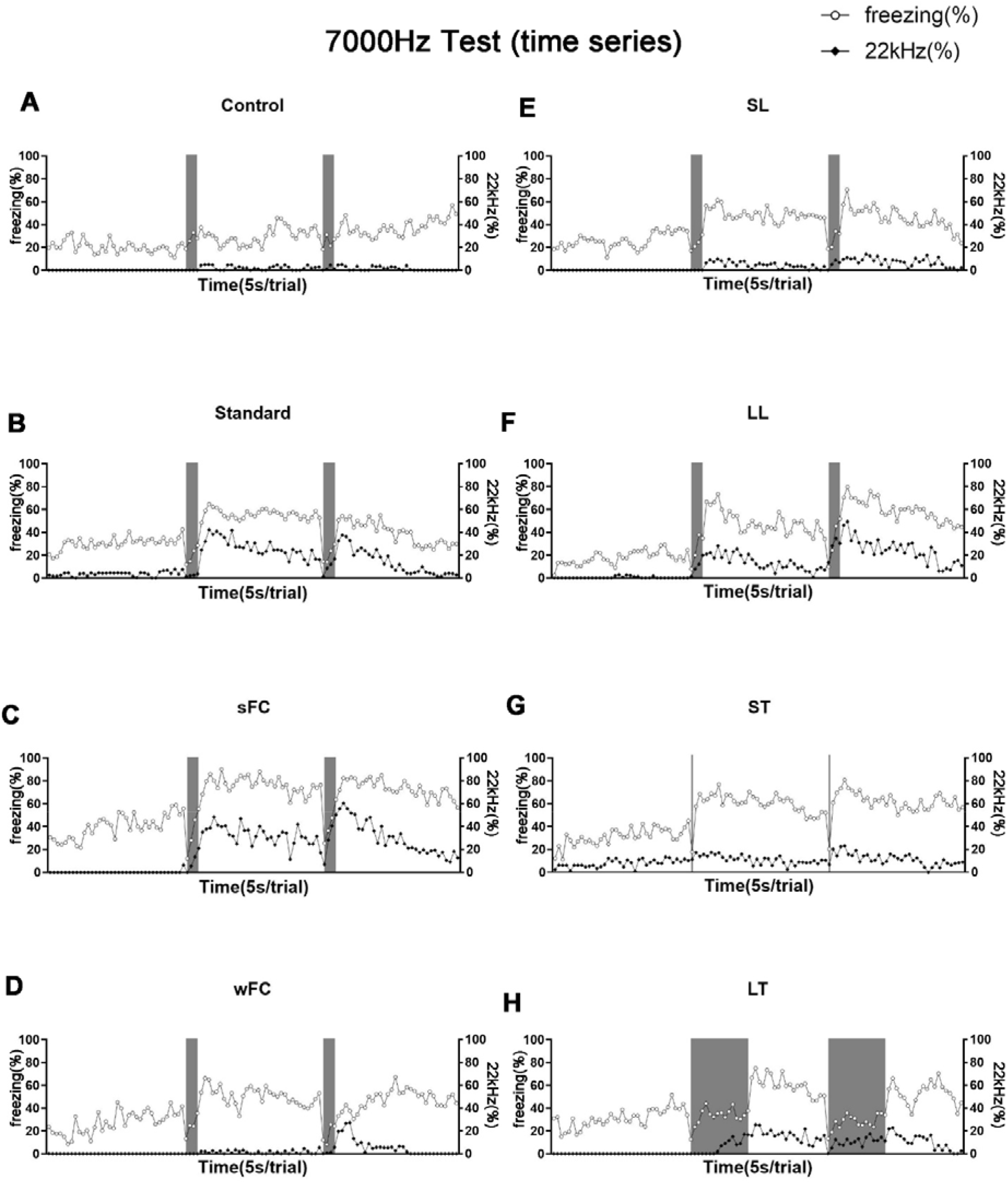
Temporal Analysis of 22kHz vocalizations and Body Freezing during 7000Hz Cue Generalization Test Across Diverse Experimental Conditions, Including the Control Group

**Figure S8:**
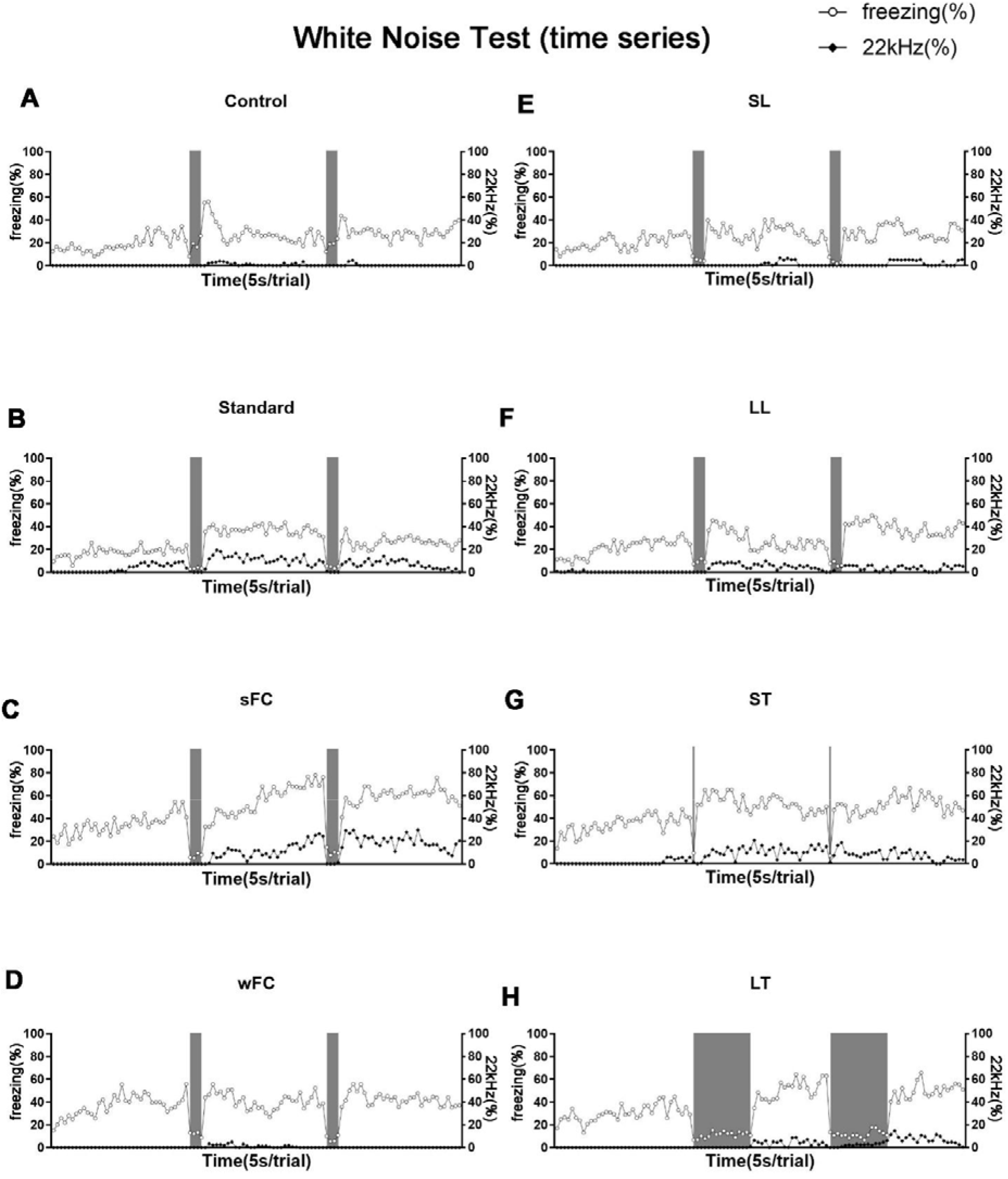
Temporal Analysis of 22kHz vocalizations and Body Freezing During White Noise Cue Generalization Test Across Diverse Experimental Conditions, Including the Control Group

**Figure S9:**
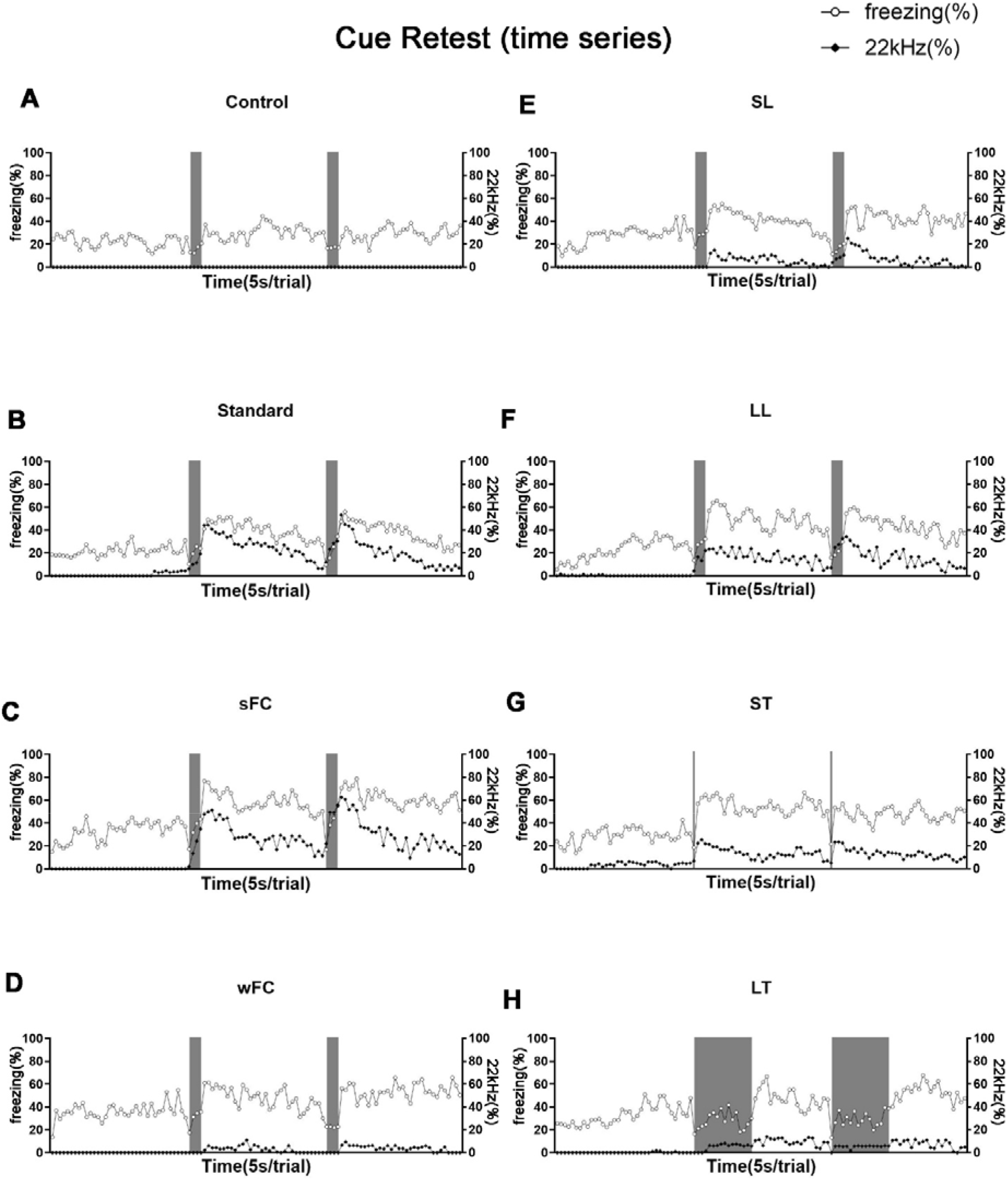
Temporal Analysis of 22kHz vocalizations and Body Freezing During 9000Hz Cue Retest Across Diverse Experimental Conditions, Including the Control Group

## Footnotes

1 We postulate that the transient increase in freezing observed in the control group can be attributed to two factors: 1. All rats had previous experiences of restraint, which potentially heightened their sensitivity to highly salient stimuli such as loud noises; 2. The volume of the CS (83-85dB) was within the threshold for highly salient stimuli. Nevertheless, the absence of 22kHz calls in the control group, despite the increase in body freezing, further substantiates our assertion that these two indicators are dissociable during the process of fear expression.

